# Proximity Labelling Identifies Proteins Associated with HSV-2 pUL21 at Early and Late Times After Infection

**DOI:** 10.1101/2025.10.16.682918

**Authors:** Safara M. Holder, Maike Bossert, Renée L. Finnen, Bruce W. Banfield

## Abstract

pUL21 is a conserved and multifunctional alphaherpesvirus tegument protein that is critical for both early and late stages in the herpes simplex virus type 2 (HSV-2) replication cycle; however, how pUL21 participates in these activities is poorly understood. To help elucidate the role of pUL21 in these various activities, we used a proximity-dependent biotin identification (BioID) approach by constructing an HSV-2 strain encoding pUL21 fused to the non-specific biotin ligase, miniTurbo (pUL21mT). Cells infected with this strain were treated with exogenous biotin at early and late times post-infection and biotinylated proteins were affinity-purified and identified by mass spectrometry. This approach enabled the identification of many viral and cellular proteins in proximity to pUL21mT at late times after infection, including those involved in cell adhesion and junction organization, as well as components of the spliceosome and the nuclear envelope. We also utilized this system to identify proteins in proximity to tegument-delivered pUL21mT immediately following viral entry, thereby providing a comprehensive profile of pUL21 proximal interactors at both early and late stages of infection. These proximal interactions were further validated by pUL21 affinity purification experiments, which confirmed that many viral and cellular proteins identified by BioID associate with pUL21. These findings provide insights into the role of HSV-2 pUL21 during infection and highlight BioID as a powerful tool for investigating virus-host interactions at multiple stages of infection.

## Introduction

Herpes simplex virus-2 (HSV-2) is a prevalent human pathogen that belongs to the *Orthoherpesviridae* family of viruses. All herpes virions share a common structure: a double-stranded DNA genome enclosed within an icosahedral capsid that is enveloped with a lipid membrane containing numerous virus-encoded envelope proteins. Located between the envelope and capsid is a compartment called the tegument that, in the case of HSV, encloses roughly 20 virus-encoded proteins and approximately 50 host cell-encoded proteins [1].

During infection of host cells, the virion envelope fuses with a cellular membrane, releasing the virion capsid into the cytoplasm. The capsid is subsequently transported along microtubules towards the nucleus, where it docks at a nuclear pore complex (NPC) and ejects the viral DNA genome into the nucleoplasm. Concomitant with the entry of capsids into the cytoplasm of the infected cell, most tegument proteins rapidly dissociate from the capsid [2–5], while others, such as those that bind dynein and kinesin motors required for capsid transport to NPCs, remain capsid-associated [3, 6–12]. With a few exceptions, the initial functions of the dispersed tegument proteins are poorly understood. Well-known examples of initial functions include the viral host shutoff (vhs) protein, a viral ribonuclease that interferes with the synthesis of host cell proteins by degrading host cell mRNAs early in infection [13, 14], and VP16 (also known as alpha-TIF and VMW65), which functions as part of a transcriptional transactivator complex to activate the expression of viral immediate early genes [15–19]. Accordingly, it is widely accepted that the tegument proteins delivered to the cell upon infection help establish a favourable environment for viral infection [20].

During virion morphogenesis, nascent nucleocapsids undergo a process termed nuclear egress to translocate from the nucleoplasm into the cytoplasm of infected cells (reviewed in [21]). This process requires that nucleocapsids undergo primary envelopment at the inner nuclear membrane (INM) as they bud into the perinuclear space. Primary envelopment of HSV nucleocapsids requires the nuclear egress complex (NEC) composed of two conserved viral proteins, pUL34 and pUL31 [21–26]. The activity of the NEC is regulated by phosphorylation, such that when the complex is hyperphosphorylated by the viral serine/threonine kinase pUs3, capsid envelopment is restricted, while when the NEC is hypophosphorylated it is active [27–31]. Following primary envelopment, primary enveloped virions fuse with the outer nuclear membrane (ONM), releasing the nucleocapsid into the cytoplasm [32]. Cytoplasmic nucleocapsids likely utilize microtubule-based transport to travel towards membranes derived from the trans-Golgi network or late endosomes. These nucleocapsids acquire their final envelope by budding into these membranes, a process termed secondary envelopment, where they become mature virions [33]. The virion tegument is acquired in stages with some tegument proteins associating with capsids in the nucleus and others being added after nuclear egress, either in the cytoplasm or through their interactions with the cytoplasmic tails of viral envelope proteins during secondary envelopment. Mature virions within vesicles are transported to and released at the plasma membrane or at membranes associated with cell junctions [33, 34]. At cell junctions, virions undergo cell-to-cell spread, enabling rapid dissemination of infection whilst circumventing detection by neutralizing antibodies [34].

pUL21 is a multifunctional tegument protein that is conserved amongst the members of the *Alphaherpesvirinae* subfamily. During the early stages of infection, it has been suggested that pUL21 promotes nucleocapsid transport along microtubules [35–37], retention of viral genomes within capsids [38], and viral gene expression [35, 39]. Throughout the later stages of infection, pUL21 regulates nuclear egress by mediating the dephosphorylation of the NEC, thereby driving primary envelopment [31, 35, 40, 41]. Graham and colleagues have established that HSV-1 pUL21 is a protein phosphatase 1 (PP1) adaptor that directs the dephosphorylation of pUs3 substrates, including the NEC [40]. While this has not yet been confirmed in HSV-2 infected cells, HSV-2 pUL21 likely mediates the dephosphorylation of the NEC through a similar mechanism. In addition to its role during nuclear egress, pUL21 prevents docking of nascent cytoplasmic nucleocapsids to NPCs [42], promotes secondary envelopment [43–45], and mediates cell-to-cell spread of infection [35, 40, 43–49]. Recently, we have demonstrated that, unlike HSV-1, HSV-2 pUL21 is phosphorylated by the viral serine/threonine kinase, pUL13, and the disruption of pUL21 phosphorylation impairs secondary envelopment and cell-to-cell spread of infection [44]. Other functions of pUL21 include its ability to facilitate the degradation of cyclic GMP-AMP synthase (cGAS) via selective autophagy [50]. Although many functions of pUL21 have been described, the mechanisms underlying most of these activities remain unclear.

To advance our understanding of the multifunctionality of pUL21 during HSV infection, we employed a proximity-dependent biotin identification (BioID) approach by constructing and characterizing an HSV-2 strain that encodes the bait protein, pUL21 fused to a non-specific biotin ligase, miniTurbo (mT; 21-mT). In the presence of exogenous biotin, mT biotinylates proteins within a 10 nm radius [51], and these biotinylated proteins can be affinity-purified and subsequently identified by mass spectrometry [52–54]. This approach enabled the identification of many viral and cellular proteins in proximity to pUL21mT at late times after infection, including those involved in cell adhesion and junction organization, as well as components of the spliceosome and the nuclear envelope. Given the robustness of this assay, this system was also utlized to identify proteins in proximity to tegument-delivered pUL21mT immediately following viral entry, thereby providing a comprehensive profile of pUL21 proximal interactors at both early and late stages of infection. These proximal interactions were further validated by pUL21 affinity purification experiments, which confirmed that 87% of viral and 42-53% of cellular proteins identified by BioID associate with pUL21. Not only do these findings provide additional insights into the role of HSV-2 pUL21 during infection, but they also highlight BioID as a powerful tool for investigating virus-host interactions at multiple stages of infection.

## Results

### Analysis of an HSV-2 strain expressing pUL21 fused to the biotin ligase miniTurbo (mT)

To enable the identification of proteins in proximity to pUL21 during infection, we constructed an HSV-2 186 strain that expresses pUL21 fused to the promiscuous biotin ligase miniTurbo (mT) [52, 53] at the UL21 locus using *en passant* mutagenesis of an HSV-2 186 BAC [35, 55]. This strain, 21-mT, replicated to a significantly lower level in human keratinocytes (HaCaT) than the wild-type (WT) 186 parental strain and at a similar level to the UL21 null strain, ΔUL21, in the absence of biotin. In the presence of biotin, the differences in replication between WT and 21-mT were not significant (Figure 1A). 21-mT formed plaques on HaCaT cells that were intermediate in size between the large WT plaques and the small plaques formed by ΔUL21 in both the presence and absence of exogenous biotin (Figure 1B, C). Taken together, these results indicate that the pUL21mT fusion protein maintains activities that support virus replication and spread, and that 21-mT is suitable for the identification of pUL21 proximal proteins.

**Figure 1.**
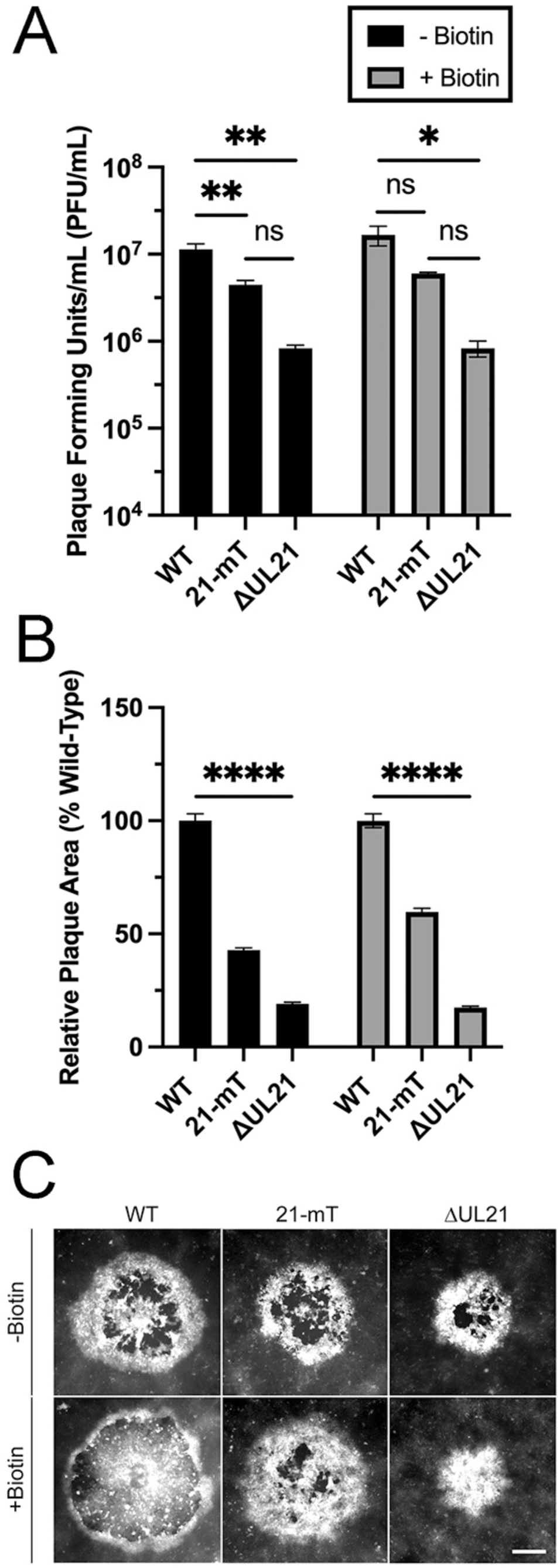
Analysis of HSV-2 pUL21mT. **(A)** The fusion of mT to pUL21 results in a virus that maintains pUL21 activity. HaCaT cells were infected in triplicate with HSV-2 strain 186 (WT), HSV-2 pUL21mT (21-mT), and HSV-2 ΔUL21 (ΔUL21) at an MOI of 0.1 in the absence or presence of exogenous biotin. At 48 hours post-infection, cells and medium were pooled, and infectious virus was titrated on HaCaT21 cells. Each data set represents the average titer from each biological triplicate. Error bars represent the standard error of the mean (*, p<0.05, **, p<0.01). **(B)** Plaque size analysis of WT, 21-mT, and ΔUL21 in the absence and presence of biotin. At 48 hours post-infection, cells were fixed, permeabilized, and stained for the viral protein, pUs3. The average plaque area of each strain relative to WT is shown (n=55 for each strain). Error bars represent the standard error of the mean (****, p<0.0001). **(C)** Representative images of plaques produced by each virus strain. The scale bar is 150 µm and applies to all panels.

### Analysis of proteins biotinylated by 21-mT during infection

HaCaT cells were infected with 21-mT and were treated with 50µM biotin for one hour at 6, 10, or 18 hours post-infection (hpi) prior to the preparation of whole cell lysates. Biotinylated proteins from cell lysates were visualized on western blots using streptavidin-conjugated horse radish peroxidase and a chemiluminescent substrate (Figure 2). As expected, endogenous biotinylated mitochondrial proteins of roughly 130 kDa (pyruvate carboxylase) and 80 kDa (propionyl-CoA carboxylase alpha chain and methylcrotonoyl-CoA carboxylase subunit alpha) were identified in mock-infected lysates and in lysates from infected cells that were not treated with biotin [56]. Many biotinylated infected cell proteins were observed in lysates prepared from 21-mT infected cells that were treated with biotin. The biotinylated protein banding pattern was similar across infection timepoints, with the most robust labelling observed at 18 hpi (Figure 2A). Biotinylated proteins were efficiently purified using streptavidin-conjugated Sepharose beads as evidenced by the lack of biotinylated proteins in the post-pull-down supernatants (Figures 2B, 2C). The subcellular localization of proteins biotinylated by pUL21mT at 18 hpi was determined by staining HaCaT cells with Alexa Fluor 488-labelled streptavidin (Figure 3). Whereas endogenous biotinylated proteins that localized to mitochondria were observed in mock-infected cells and in 21-mT infected cells that were not treated with biotin (arrows), robust pancellular labelling of biotinylated proteins was observed in biotin-treated 21-mT infected cells. Consistent with the reported localization of pUL21 to the nuclear rim, biotin labelling of the nuclear envelope was readily apparent in 21-mT infected cells (arrowheads) [35, 40, 42].

**Figure 2.**
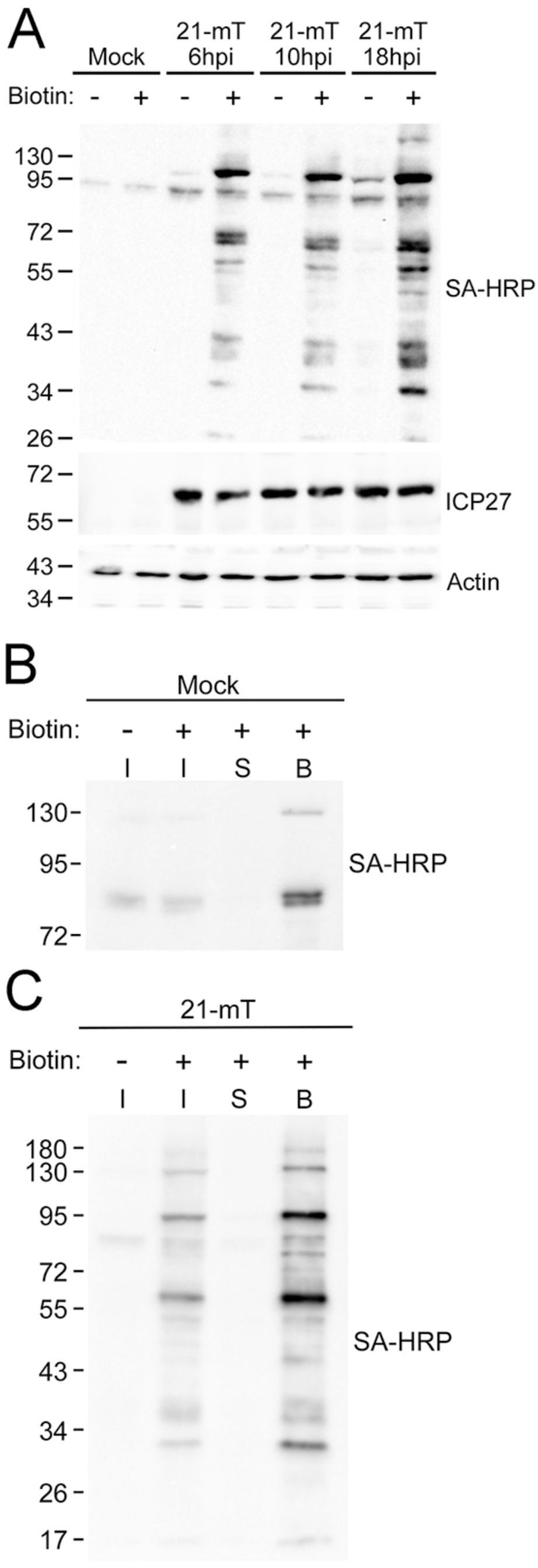
The biotinylation profile of HSV-2 pUL21mT in HaCaT cells. **(A)** Time course of pUL21mT biotinylation. HaCaT cells were mock-infected or infected with HSV-2 pUL21mT (21-mT) at an MOI of 1 and treated with exogenous biotin for one hour at 6,10, and 18 hours post-infection (hpi). Biotinylated proteins were visualized by western blotting using streptavidin-HRP (SA-HRP). Antibodies against actin and the viral protein ICP27 were used as loading and infectivity controls, respectively. **(B & C)** Affinity purification of proteins biotinylated by pUL21mT at 18 hpi. HaCaT cells were mock-infected **(B)** or infected with 21-mT **(C)** for 18 hours and treated with exogenous biotin for one hour. Biotinylated proteins were affinity-purified using streptavidin-Sepharose beads. Following incubation, bead-bound proteins were pelleted by centrifugation (Bound; B), and post-pull-down supernatants (S) were collected. A portion of each sample was retained prior to the addition of streptavidin-Sepharose beads (Input; I). Biotinylated proteins were visualized by western blotting using SA-HRP. Migration positions of molecular weight markers (kDa) are shown on the left side of each western blot.

**Figure 3.**
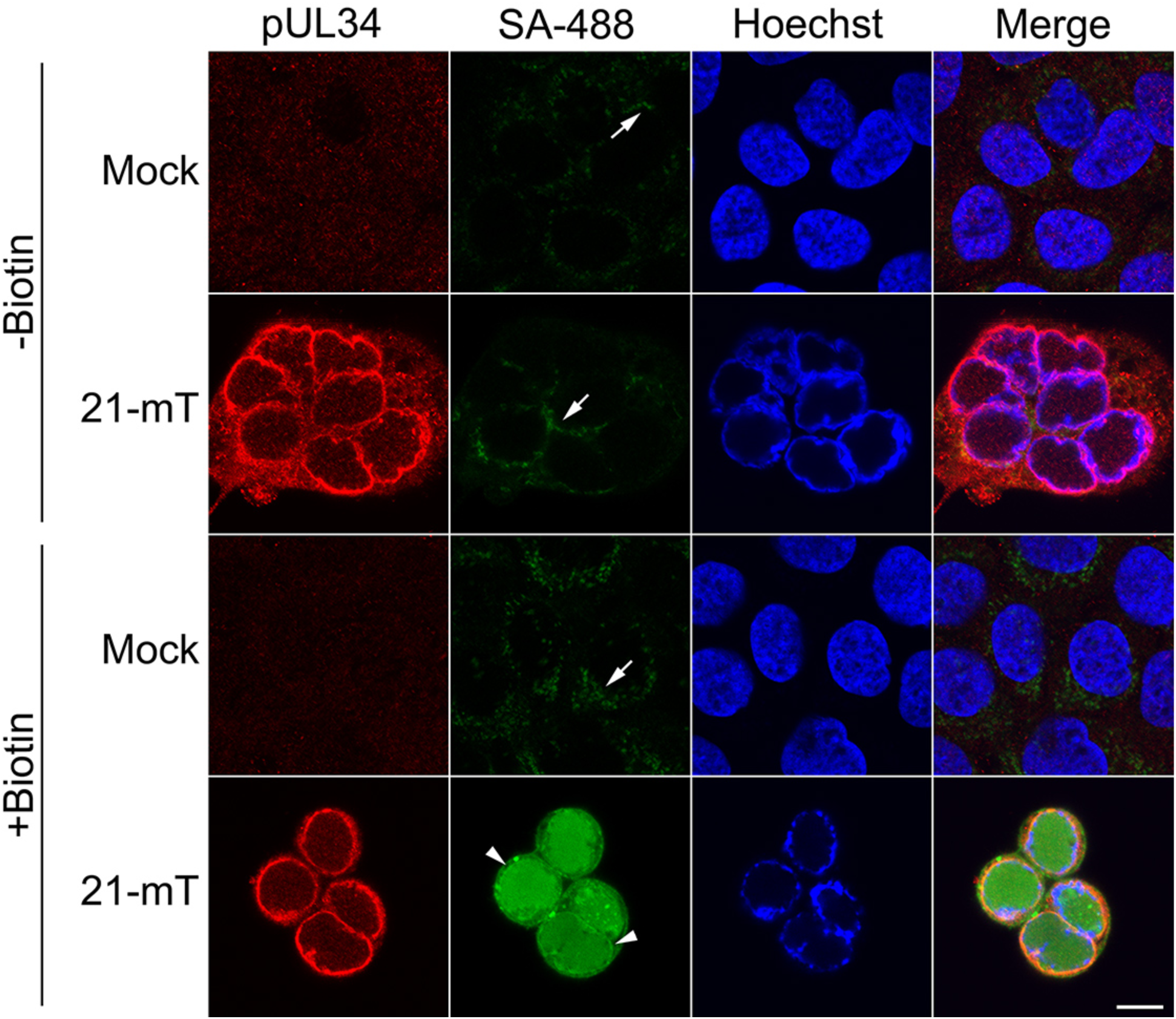
Subcellular localization of proteins biotinylated by pUL21mT at 18 hours post-infection. HaCaT cells were mock-infected or infected with HSV-2 pUL21mT (21-mT) at an MOI of 0.3. At 18 hours post-infection, cells were treated with exogenous biotin for one hour, fixed, permeabilized, and stained with antisera against the viral protein pUL34 (red), streptavidin-conjugated Alexa Fluor 488 (SA-488; green), and the DNA stain Hoechst 33342 (blue). Arrows and arrowheads point to mitochondria containing endogenous biotinylated proteins and the nuclear envelope, respectively. The scale bar is 10 µm and applies to all panels.

### Identification of proteins biotinylated by 21-mT at 18 hpi

HaCaT cells were infected in triplicate with 21-mT for 18 hours, treated with or without 50µM biotin for one hour, after which biotinylated proteins were affinity-purified on streptavidin-conjugated Sepharose beads. Proteins on beads were digested with trypsin and resulting peptides were identified by LC-MS/MS. A total of 640 cellular and viral proteins were identified in this analysis (supplemental data 1). These were further distilled by excluding proteins that: 1) were common contaminants; 2) had less than an average of three peptides identified in each of the three biological replicates; and 3) had a normalized percent coverage less than 5%. Based on these criteria, 110 cellular and 13 viral proteins were further analyzed. Table 1 lists the top 50 cellular proteins in proximity to pUL21mT ranked by normalized percent coverage and Table 2 lists the top 13 viral proteins in proximity to pUL21mT, ranked by normalized percent coverage.

**Table 1.** Top 50 Cellular Proteins Identified in Proximity to pUL21mT at 18 hpi.

| <sup>1</sup>Rank | Protein Name | Gene Name | Molecular Weight | <sup>2</sup>Normalized Spectral Counts | <sup>3</sup>Normalized Percent Coverage <sup>4</sup>(±SD) |
| --- | --- | --- | --- | --- | --- |
| 1 | Lamina-associated polypeptide 2, isoforms beta/gamma | TMPO | 51 kDa | 71.2 | 53.1 (±1.7) |
| 2 | Cytoplasmic dynein 1 light intermediate chain 2 | DC1L2_HUMAN | 54 kDa | 27.9 | 44.5 (±6) |
| 3 | Transgelin-2 | TAGL2_HUMAN | 22 kDa | 11.6 | 41 (±5.5) |
| 4 | Annexin A1 | ANXA1_HUMAN | 39 kDa | 20.5 | 40 (±1.8) |
| 5 | <sup>5</sup>Serine/threonine-protein phosphatase PP1-alpha catalytic subunit | PP1A_HUMAN | 38 kDa | 20.8 | 36.7 (±3.1) |
| 6 | T-complex protein 1 subunit theta | TCPQ_HUMAN | 60 kDa | 33.4 | 35.8 (±1.5) |
| 7 | Plakophilin-3 | PKP3_HUMAN | 87 kDa | 34.6 | 35.5 (±2.5) |
| 8 | Serine/threonine-protein phosphatase PP1-gamma catalytic subunit | PP1G_HUMAN | 37 kDa | 19.7 | 34.9 (±3.4) |
| 9 | Heterogeneous nuclear<br>ribonucleoprotein A1 | ROA1_HUMAN | 39 kDa | 54.8 | 30.6 ($\pm 2$) |
| 10 | Cytoplasmic dynein 1 light<br>intermediate chain 1 | DC1L1_HUMAN | 57 kDa | 25.3 | 30.5 ($\pm 5.4$) |
| 11 | Protein LYRIC | LYRIC_HUMAN | 64 kDa | 22.8 | 29.8 ($\pm 5.7$) |
| 12 | Signal recognition particle receptor<br>subunit alpha | SRPRA_HUMAN | 70 kDa | 27 | 29.8 ($\pm 3.5$) |
| 13 | Inner nuclear membrane protein<br>Man1 | MAN1_HUMAN | 100 kDa | 35.3 | 29.7 ($\pm 2.2$) |
| 14 | Unconventional myosin-VI | MYO6_HUMAN | 150 kDa | 66.1 | 29.3 ($\pm 3$) |
| 15 | U4/U6 small nuclear<br>ribonucleoprotein Prp3 | PRPF3_HUMAN | 78 kDa | 27.1 | 28.7($\pm 2.1$) |
| 16 | Tripartite motif-containing protein 29 | TRI29_HUMAN | 66 kDa | 26.7 | 28.6 ($\pm 4.4$) |
| 17 | Pre-mRNA-splicing factor SYF1 | SYF1_HUMAN | 100 kDa | 39.2 | 28.5 ($\pm 1.3$) |
| 18 | Stromal interaction molecule 1 | STIM1_HUMAN | 77 kDa | 32.7 | 28.4 ($\pm 2.9$) |
| 19 | Transcription factor jun-B | JUNB_HUMAN | 36 kDa | 10 | 28 ($\pm 0$) |
| 20 | Synaptotagmin-like protein 4 | SYTL4_HUMAN | 76 kDa | 30.3 | 27.6 ($\pm 3.4$) |
| 21 | Beta/gamma crystallin domain-containing protein 1 | CRBG1_HUMAN | 189 kDa | 63.4 | 27.4 ($\pm 0.6$) |
| 22 | Heterogeneous nuclear<br>ribonucleoprotein K | HNRPK_HUMAN | 51 kDa | 20.6 | 26.7(±2.4) |
| 23 | Src substrate cortactin | SRC8_HUMAN | 62 kDa | 20.4 | 25.1 (±2.2) |
| 24 | Heterogeneous nuclear<br>ribonucleoproteins A2/B1 | ROA2_HUMAN | 37 kDa | 17 | 23.8 (±3.5) |
| 25 | Proliferation marker protein Ki-67 | KI67_HUMAN | 359 kDa | 115.6 | 23.8 (±3.6) |
| 26 | Delta(14)-sterol reductase LBR | LBR_HUMAN | 71 kDa | 30.2 | 22.8 (±1.2) |
| 27 | SUN domain-containing protein 1 | SUN1_HUMAN | 90 kDa | 23.2 | 22.5 (±3.4) |
| 28 | Myristoylated alanine-rich C-kinase<br>substrate | MARCS_HUMAN | 32 kDa | 7.4 | 22.3(±2.6) |
| 29 | Nuclear pore complex protein<br>Nup153 | NU153_HUMAN | 154 kDa | 41.8 | 21.9 (±2.5) |
| 30 | Torsin-1A-interacting protein 1 | TOIP1_HUMAN | 66 kDa | 17.1 | 21.6 (±2.3) |
| 31 | Epidermal growth factor receptor<br>substrate 15-like 1 | EP15R_HUMAN | 94 kDa | 22.6 | 20.7 (±2.8) |
| 32 | Probable ATP-dependent RNA<br>helicase DDX5 | DDX5_HUMAN | 69 kDa | 23.5 | 20.4 (±2.4) |
| 33 | Protein ELYS | ELYS_HUMAN | 253 kDa | 62.2 | 19.3 (±2.5) |
| 34 | Transcriptional coactivator YAP1 | YAP1_HUMAN | 54 kDa | 11 | 19.3 (±0.8) |
| 35 | E3 SUMO-protein ligase RanBP2 | RBP2_HUMAN | 358 kDa | 78.7 | 18.7 (± 1.5) |
| 36 | Cell cycle and apoptosis regulator<br>protein 2 | CCAR2_HUMAN | 103 kDa | 18.5 | 18.2 (±2.8) |
| 37 | Emerin | EMD_HUMAN | 29 kDa | 7 | 18 (±1.2) |
| 38 | Heterogeneous nuclear<br>ribonucleoprotein A3 | ROA3_HUMAN | 40 kDa | 8.9 | 17.7 (±2.1) |
| 39 | Poly(rC)-binding protein 1 | PCBP1_HUMAN | 37 kDa | 7.5 | 17.7 (±0) |
| 40 | Splicing factor 3B subunit 1 | SF3B1_HUMAN | 146 kDa | 28.8 | 17.5 (±3.9) |
| 41 | Arf-GAP domain and FG repeat-containing protein 1 | AGFG1_HUMAN | 58 kDa | 17.9 | 17.4 (±0) |
| 42 | Lymphoid-specific helicase | HELLS_HUMAN | 97 kDa | 26 | 17.1 (±1.9) |
| 43 | Endoplasmic reticulum chaperone<br>BiP | BIP_HUMAN | 72 kDa | 12.6 | 16.9 (±3.2) |
| 44 | ATP-dependent RNA helicase<br>DDX3X | DDX3X_HUMAN | 73 kDa | 13.7 | 16.8 (±6.3) |
| 45 | Prelamin-A/C | LMNA_HUMAN | 74 kDa | 20.1 | 16.7 (±4.4) |
| 46 | Epidermal growth factor receptor | EGFR_HUMAN | 134 kDa | 25.1 | 16.3 (±1.4) |
| 47 | U4/U6.U5 tri-snRNP-associated<br>protein 1 | SNUT1_HUMAN | 90 kDa | 15.7 | 16.3 (±3.5) |
| 48 | Long-chain-fatty-acid--CoA ligase 4 | ACSL4_HUMAN | 79 kDa | 14 | 16.2 (±1.3) |
| 49 | BUD13 homolog | BUD13_HUMAN | 71 kDa | 14.6 | 16.1 (±0.8) |
| 50 | Gamma-interferon-inducible protein<br>16 | IF16_HUMAN | 88 kDa | 49.3 | 16 (±2.2) |

<sup>1</sup>Rank based on normalized percent coverage.
<sup>2</sup>Spectral counts from three biological replicates were normalized to endogenously biotinylated cellular proteins then averaged. Averages of the no-biotin control samples were subtracted from the plus-biotin experimental samples to determine the normalized spectral count.
<sup>3</sup>Average of the percent coverage of three biological replicates of the no-biotin control samples was subtracted from the average of the percent coverage of the plus-biotin experimental samples to determine the normalized percent coverage.
<sup>4</sup>Standard deviation (SD) of the three plus-biotin biological replicates.
<sup>5</sup>Peptides common to both PP1A\_HUMAN and PP1G\_HUMAN were identified as were peptides unique to each protein.

**Table 2.**
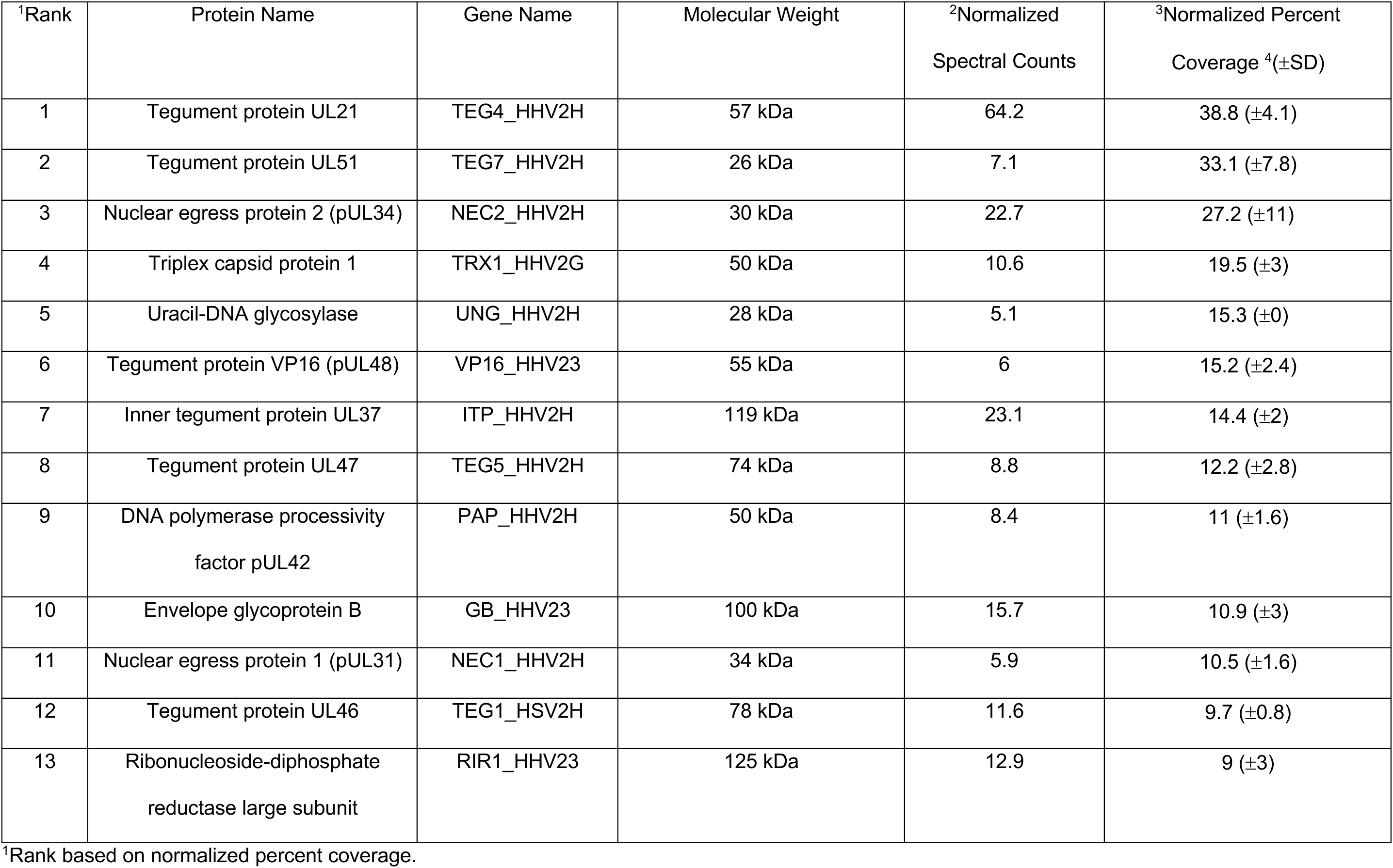
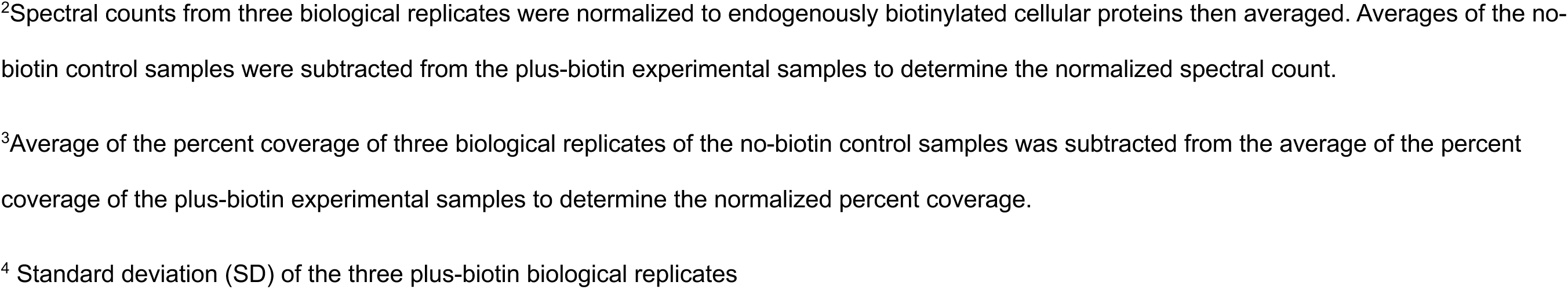
Top 13 Viral Proteins Identified in Proximity to pUL21mT at 18 hpi.

| <sup>1</sup>Rank | Protein Name | Gene Name | Molecular Weight | <sup>2</sup>Normalized Spectral Counts | <sup>3</sup>Normalized Percent Coverage <sup>4</sup>(±SD) |
| --- | --- | --- | --- | --- | --- |
| 1 | Tegument protein UL21 | TEG4_HHV2H | 57 kDa | 64.2 | 38.8 (±4.1) |
| 2 | Tegument protein UL51 | TEG7_HHV2H | 26 kDa | 7.1 | 33.1 (±7.8) |
| 3 | Nuclear egress protein 2 (pUL34) | NEC2_HHV2H | 30 kDa | 22.7 | 27.2 (±11) |
| 4 | Triplex capsid protein 1 | TRX1_HHV2G | 50 kDa | 10.6 | 19.5 (±3) |
| 5 | Uracil-DNA glycosylase | UNG_HHV2H | 28 kDa | 5.1 | 15.3 (±0) |
| 6 | Tegument protein VP16 (pUL48) | VP16_HHV23 | 55 kDa | 6 | 15.2 (±2.4) |
| 7 | Inner tegument protein UL37 | ITP_HHV2H | 119 kDa | 23.1 | 14.4 (±2) |
| 8 | Tegument protein UL47 | TEG5_HHV2H | 74 kDa | 8.8 | 12.2 (±2.8) |
| 9 | DNA polymerase processivity factor pUL42 | PAP_HHV2H | 50 kDa | 8.4 | 11 (±1.6) |
| 10 | Envelope glycoprotein B | GB_HHV23 | 100 kDa | 15.7 | 10.9 (±3) |
| 11 | Nuclear egress protein 1 (pUL31) | NEC1_HHV2H | 34 kDa | 5.9 | 10.5 (±1.6) |
| 12 | Tegument protein UL46 | TEG1_HSV2H | 78 kDa | 11.6 | 9.7 (±0.8) |
| 13 | Ribonucleoside-diphosphate reductase large subunit | RIR1_HHV23 | 125 kDa | 12.9 | 9 (±3) |

<sup>1</sup>Rank based on normalized percent coverage.
<sup>2</sup>Spectral counts from three biological replicates were normalized to endogenously biotinylated cellular proteins then averaged. Averages of the no- biotin control samples were subtracted from the plus-biotin experimental samples to determine the normalized spectral count.
<sup>3</sup>Average of the percent coverage of three biological replicates of the no-biotin control samples was subtracted from the average of the percent coverage of the plus-biotin experimental samples to determine the normalized percent coverage.
<sup>4</sup> Standard deviation (SD) of the three plus-biotin biological replicates

Gene ontogeny analysis of cellular proteins was performed using Metascape [57, 58] and is shown in Figure 4A. Many proteins involved in cadherin binding, mRNA metabolism, and nuclear envelope breakdown were identified. STRING 12.0 analysis [57, 59, 60] of protein-protein interaction networks of the cellular proteins identified 7 main interaction clusters, including cell junction organization, the spliceosome, nuclear envelope breakdown, mRNA processing and stability, and the clathrin coat (Figure 4B).

**Figure 4.**
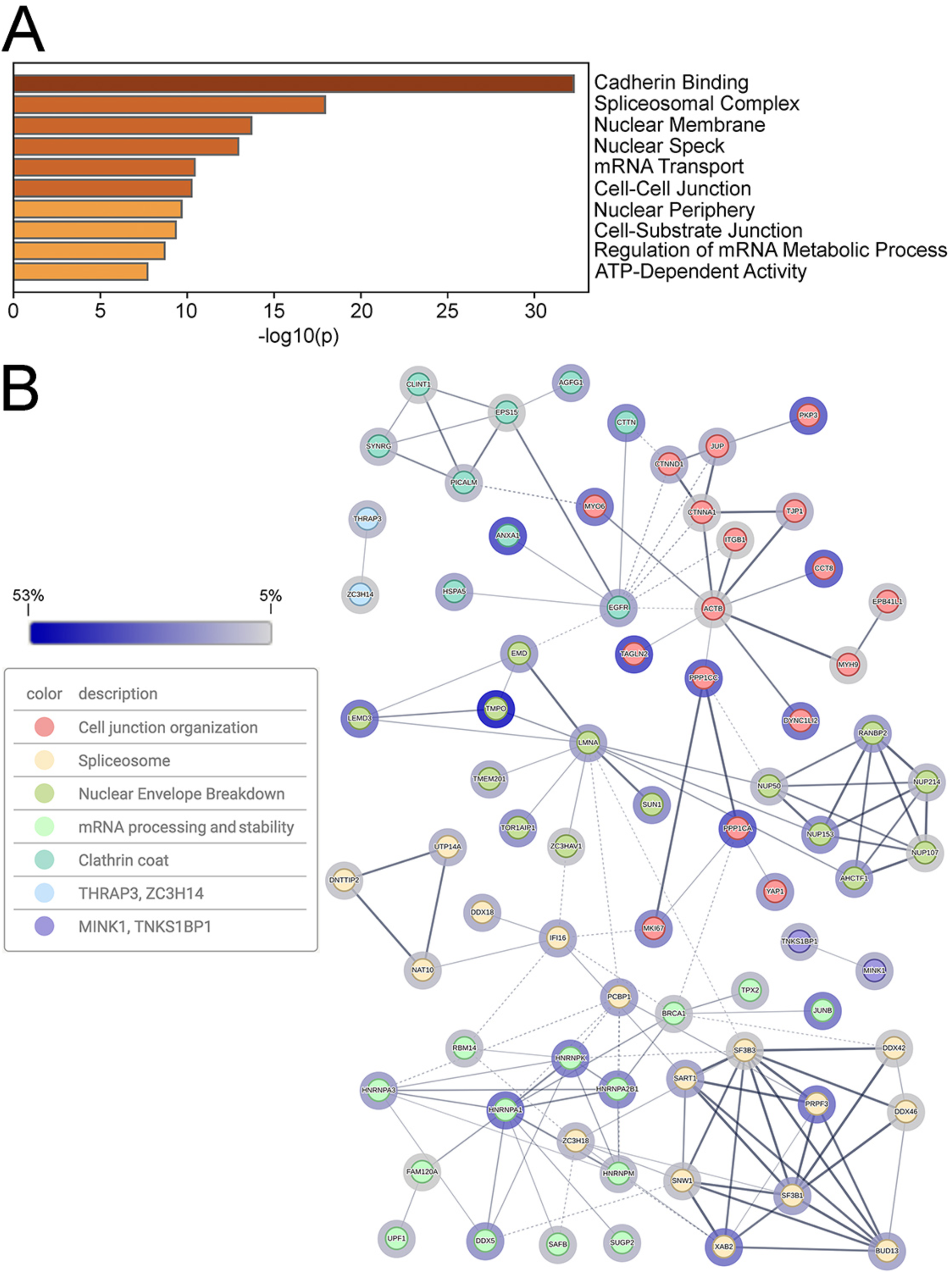
The identification of proteins in proximity to pUL21mT at 18 hours post-infection. **(A)** Top 10 Metascape-generated gene ontologies of cellular proteins identified in proximity to pUL21mT at 18 hours post-infection. **(B)** STRING network of cellular proteins in proximity to pUL21mT. Identified proteins were analyzed, clustered, and colour-coded using the STRING database to visualize known protein-protein interactions. Edges connecting the nodes represent reported associations, with edge thickness corresponding to interaction confidence (thicker edges indicate higher confidence). Halos surrounding each node reflect percent coverage, with darker halos indicating higher percent coverage. Only proteins with a normalized protein coverage of ≥ 5% were included in this analysis, and disconnected nodes are not shown.

Given the role of pUL21 in nuclear egress [31, 35, 40, 41], it was intriguing to see that in addition to the lamina component prelamin A/C, numerous cellular and viral INM proteins were in proximity to pUL21mT at 18 hpi. These included the cellular proteins lamina-associated polypeptide 2, inner nuclear membrane protein Man1, delta (14)-sterol reductase lamin B receptor (LBR), SUN domain-containing protein 1, torsin-1A-interacting protein 1, emerin, and the viral NEC proteins pUL34 and pUL31. Considering that pUL21 lacks a transmembrane domain or any canonical signals that might recruit it to the nuclear lamina or nuclear envelope, it is likely that one or more of these proteins are involved in the recruitment of pUL21 to the nuclear rim. To further evaluate the biophysical properties of pUL21 at the nuclear envelope, fluorescence recovery after photobleaching (FRAP) analysis was performed. Consistent with the behaviour of soluble nucleoplasmic proteins, FRAP analysis of HeLa cells expressing the soluble nucleoplasmic protein pUL31-EGFP showed near instantaneous recovery of fluorescence to a region of interest that had been photobleached by a 405 nm laser (Figure 5). Co-transfection of cells with a plasmid expressing the INM binding partner of pUL31, FLAG-pUL34, along with the pUL31-EGFP expression plasmid resulted in the recruitment of pUL31-EGFP to the INM. Photobleaching of INM localized pUL31-EGFP resulted in maximal recovery of fluorescence to the bleached region in approximately 100 seconds. By contrast, photobleaching of cells transfected with a plasmid encoding an EGFP fusion to the lamina component, lamin B1, resulted in no recovery of fluorescence to the photobleached region. These findings reflect the stark differences in the relative mobilities of INM proteins compared to components of the lamina. In cells transfected with a pUL21-EGFP expression plasmid, pUL21-EGFP localized to the nuclear envelope, the nucleoplasm, and the cytoplasm. These findings indicate that pUL21-EGFP recruitment to the nuclear envelope does not require the expression of additional viral proteins. Photobleaching of pUL21-EGFP at the nuclear envelope resulted in recovery of fluorescence to the bleached region with similar kinetics observed for pUL31-EGFP when recruited to the INM by FLAG-pUL34. Taken together, these findings are consistent with the recruitment of pUL21 to the nuclear envelope through interactions with cellular nuclear envelope proteins; perhaps one or more of the proteins identified in this BioID analysis.

**Figure 5.**
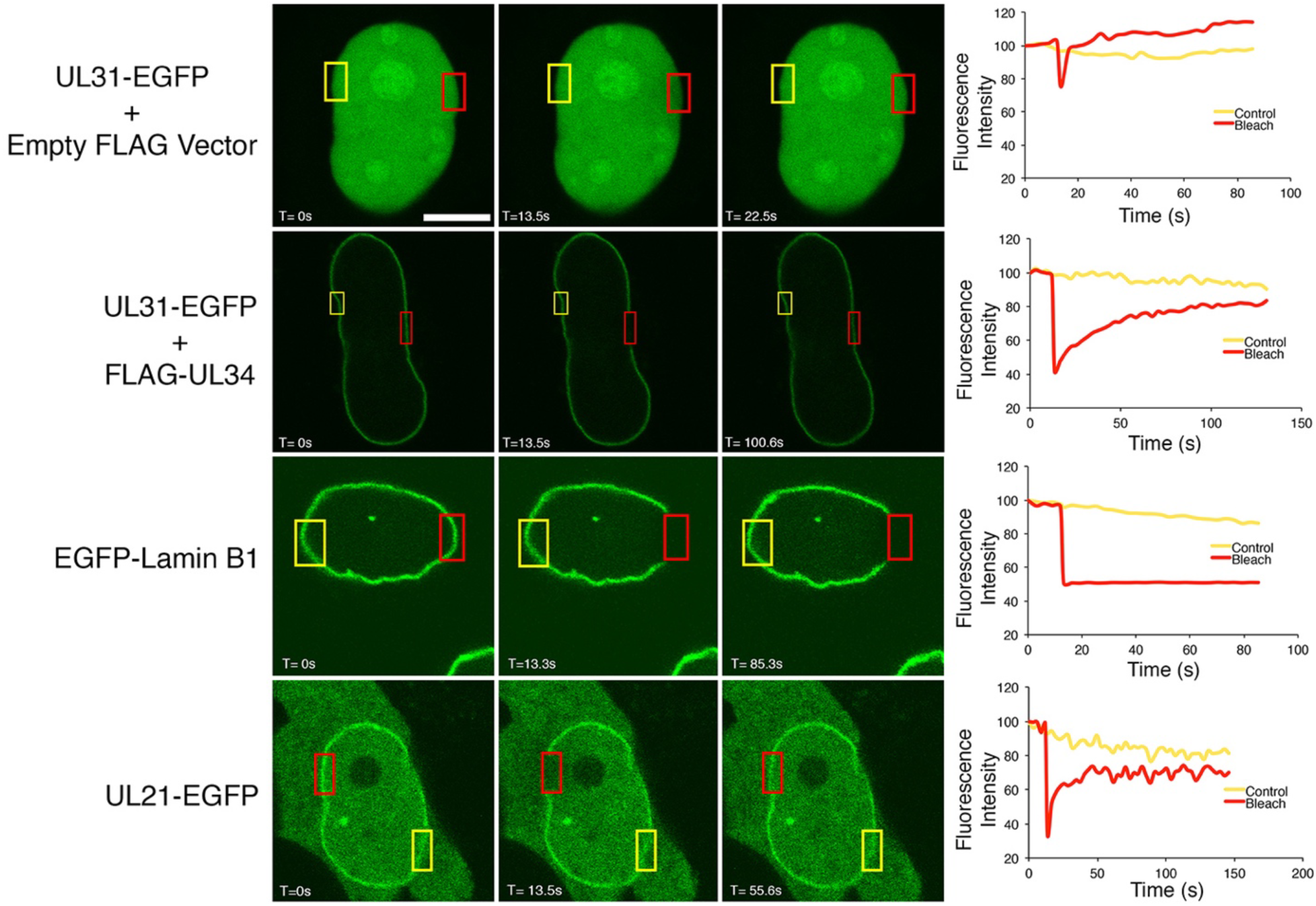
The recruitment of pUL21 to the nuclear envelope. HeLa cells were co-transfected with the indicated expression plasmids, and images of live cells were collected at 24 hours post-transfection. Regions outlined in red were photobleached, and representative images of cells immediately before, immediately after, and during the recovery period are shown. Distal non-photobleached control regions are outlined in yellow. The scale bar in the top left panel is 10 µm and applies to all images. Corresponding graphs display the fluorescence intensity of the control and photobleached regions over time.

### Analysis of proteins biotinylated by 21-mT immediately following virus entry

Because pUL21 is incorporated into the tegument of WT virions, we explored the possibility that 21-mT would package pUL21mT into the tegument of 21-mT virions. If this were the case, tegument-delivered pUL21mT might be capable of biotinylating proximal proteins immediately after virus entry. Western blot analysis of virions purified from WT and 21-mT infected cells was normalized based on levels of the major capsid protein, VP5, found in 955 copies per capsid [61]. pUL21mT was incorporated into 21-mT virions, albeit to a lesser extent than pUL21 was incorporated into WT virions (Figure 6A). To test if tegument-localized pUL21mT was capable of biotinylating virion components, purified 21-mT virions were incubated with 50 µM biotin in the presence and absence of low concentrations of saponin or NP-40 to permeabilize the virion envelope (Figure 6B). The levels of protein biotinylation in untreated 21-mT virions were modest, with three major species observed and a predominant band migrating at approximately 95 kDa, corresponding to the size of pUL21mT. Incubation of 21-mT virions with biotin in the presence of saponin did not alter the biotinylation profile, whereas NP-40 treatment resulted in a modest increase in biotinylation and the concomitant appearance of a few additional biotinylated species. These data suggest that, in the context of purified virions, the ability of pUL21mT to biotinylate virion components is limited.

**Figure 6.**
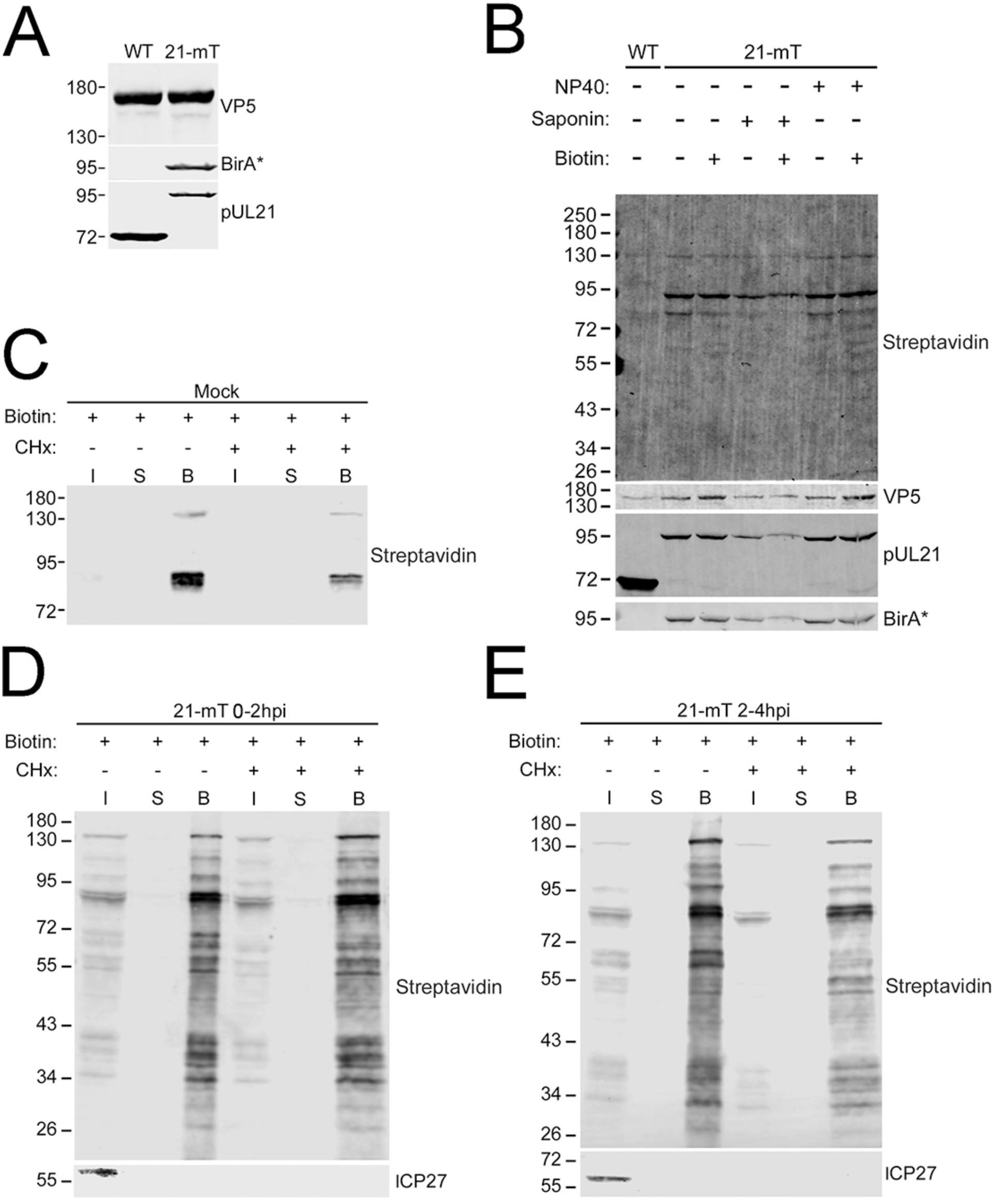
Tegument-delivered pUL21mT biotinylates infected cell proteins immediately following viral entry. **(A)** pUL21mT is packaged into HSV-2 pUL21mT virions. HSV-2 strain 186 (WT) and HSV-2 pUL21mT (21-mT) virions were purified by ultracentrifugation through a 30% sucrose cushion. Isolated virions were analyzed by western blotting using antisera against the major capsid protein, VP5, BirA* (reactive against mT) and pUL21. **(B)** pUL21mT weakly biotinylates virion components within purified 21-mT virions. HaCaT cells were infected with WT or 21-mT at an MOI of 0.01. At 72 hours post-infection, extracellular virions were purified by ultracentrifugation through a 30% sucrose cushion. Isolated virions were treated with saponin or NP40 to permeabilize the virion envelope, treated with exogenous biotin for one hour, and analyzed by western blotting using streptavidin-IRDye 680RD (Streptavidin) to visualize biotinylated proteins, antisera against the major capsid protein, VP5, pUL21, and BirA* (reactive against mT). **(C-E)** The biotinylation profile of 21-mT infected cells immediately following viral entry. HaCaT cells were mock-infected (**C**) or infected with 21-mT in the presence of cycloheximide (CHx). Cells were then treated with exogenous biotin between 0-2 (**D**) or 2-4 hours post-infection (hpi) (**E**), harvested, and incubated with streptavidin-Sepharose beads. Biotinylated proteins bound to streptavidin-Sepharose beads were pelleted by centrifugation (Bound; B), and post-pull-down supernatants (S) were collected. A portion of each sample was collected prior to the addition of streptavidin-Sepharose beads (input;I). Biotinylated proteins were visualized by western blotting using streptavidin, and antisera against the viral protein ICP27 served as a CHx and infectivity control. Migration positions of molecular weight markers (kDa) are shown on the left side of each western blot.

We next explored the possibility that tegument-delivered pUL21mT could biotinylate proximal proteins immediately after virus entry. To do this, we infected HaCaT cells with 21-mT in the presence of 50 µM biotin at 0-2 or 2-4 hpi. To ensure that any biotinylation observed was due to incoming pUL21mT rather than *de novo* synthesized pUL21mT, cells were also treated with the protein synthesis inhibitor cycloheximide (CHx). After biotinylation was complete, biotinylated proteins were purified using streptavidin-conjugated Sepharose beads, after which bound and unbound proteins were visualized by western blotting using streptavidin-conjugated HRP. In mock-infected cells, only the endogenously biotinylated mitochondrial proteins were affinity-purified (Figure 6C). When biotin was added between 0 and 2 hpi (Figure 6D), or 2 and 4 hpi (Figure 6E), many biotinylated protein species were efficiently affinity-purified. To verify that the CHx treatment prevented *de novo* protein expression, lysates from infected cells were probed for the HSV-2 immediate early protein ICP27. Whereas ICP27 was readily detected in untreated cell lysates, ICP27 was not observed in cell lysates that had been exposed to CHx. Biotinylation of proteins by tegument-delivered pUL21mT was more efficient when cells were treated with biotin between 0 and 2 hpi than between 2 and 4 hpi; thus, the 0 to 2 hpi labelling protocol was used to identify biotinylated proteins.

To determine the subcellular localization of proteins in proximity to tegument-delivered pUL21mT, 21-mT infected HaCaT cells that were incubated with biotin between 0 and 2 hpi were fixed and stained with Alexa Fluor 488-labelled streptavidin (Figure 7). Endogenous biotinylated proteins that localized to mitochondria were observed in control 21-mT infected cells that were not treated with biotin. Irrespective of the presence or absence of CHx, biotinylated proteins localized predominantly to the nucleus and the nuclear rim (arrows) in biotin-labelled infected cells.

**Figure 7.**
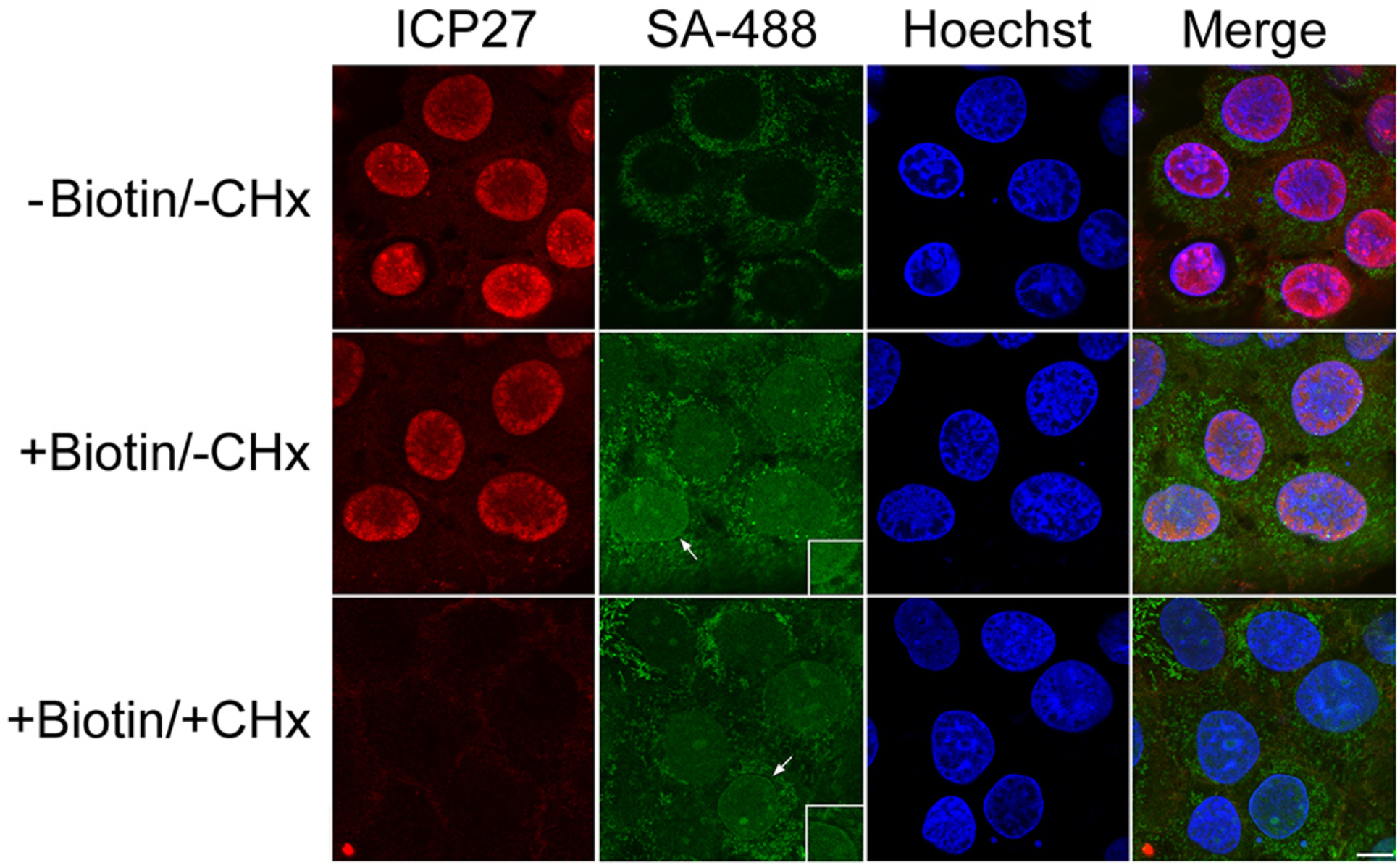
Subcellular localization of proteins biotinylated by tegument-delivered pUL21mT at two hours post-infection. HaCaT cells were infected with HSV-2 pUL21mT for two hours in the presence of exogenous biotin and cycloheximide (CHx). Cells were fixed and stained with antisera against the viral protein, ICP27 (red), streptavidin conjugated to Alexa Fluor 488 (SA-488; green) to detect biotinylated proteins, and the DNA stain Hoechst 33342 (blue). Arrows point to the nuclear rim highlighted in the zoomed inset panels. The scale bar is 10 µm and applies to all panels.

### Identification of proteins biotinylated by 21-mT at 2 hpi

CHx-treated HaCaT cells were infected in triplicate with 21-mT for 2 hours in the continuous presence, or absence, of 50 µM biotin. Biotinylated proteins were affinity-purified on streptavidin-conjugated Sepharose beads. Proteins on beads were digested with trypsin and resulting peptides were identified by LC-MS/MS. A total of 357 cellular and viral proteins were identified in this analysis (supplemental data 2). These were further distilled, as described above for the 18 hpi dataset. Based on these criteria, 53 cellular proteins were further analyzed. Interestingly, the only viral protein that fulfilled our criteria was pUL21. Table 3 lists the top 50 cellular proteins in proximity to pUL21mT at 2 hpi, ranked by normalized percent coverage.

**Table 3.** Top 50 Cellular Proteins Identified in Proximity to pUL21mT at 2 hpi.

| <sup>1</sup>Rank | Protein Name | Gene Name | Molecular Weight | <sup>2</sup>Normalized Spectral Counts | <sup>3</sup>Normalized Percent Coverage <sup>4</sup>(±SD) |
| --- | --- | --- | --- | --- | --- |
| 1 | Lamina-associated polypeptide 2, isoforms beta/gamma | TMPO | 51 kDa | 40.6 | 57.7 (±1) |
| 2 | Gamma-interferon-inducible protein 16 | IF16_HUMAN | 88 kDa | 102.3 | 50.3 (±2.5) |
| 3 | T-complex protein 1 subunit theta | TCPQ_HUMAN | 60 kDa | 20.5 | 36.9 (±1.1) |
| 4 | Unconventional myosin-VI | MYO6_HUMAN | 150 kDa | 57 | 33.9 (±0) |
| 5 | Tripartite motif-containing protein 29 | TRI29_HUMAN | 66 kDa | 23.3 | 31.5 (±0.8) |
| 6 | Thioredoxin | THIO_HUMAN | 12 kDa | 7.2 | 31.4 (±0.5) |
| 7 | Synaptotagmin-like protein 4 | SYTL4_HUMAN | 76 kDa | 23.2 | 30.3 (±0) |
| 8 | Erbin | ERBIN_HUMAN | 158 kDa | 37.8 | 29.3 (±7.8) |
| 9 | Junction plakoglobin | PLAK_HUMAN | 82 kDa | 25.4 | 28.6 (±6) |
| 10 | Pleckstrin homology domain-containing family A member 5 | PKHA5_HUMAN | 127 kDa | 33.8 | 28.1 (±0.7) |
| 11 | Catenin delta-1 | CTND1_HUMAN | 108 kDa | 34.7 | 26.8 (±4.7) |
| 12 | <sup>5</sup>Serine/threonine-protein phosphatase PP1-gamma catalytic subunit | PP1G_HUMAN | 37 kDa | 8.5 | 26.3 (±11.5) |
| 13 | Inner nuclear membrane protein Man1 | MAN1_HUMAN | 100 kDa | 15.7 | 23.7 ($\pm 0$) |
| 14 | Heterogeneous nuclear<br>ribonucleoprotein A1 | ROA1_HUMAN | 39 kDa | 10.2 | 23.1 ($\pm 5.2$) |
| 15 | Emerin | EMD_HUMAN | 29 kDa | 5.1 | 22.6 ($\pm 0.2$) |
| 16 | Serine/threonine-protein phosphatase PP1-alpha catalytic subunit | PP1A_HUMAN | 38 kDa | 6.4 | 21.9 ($\pm 1.1$) |
| 17 | Src substrate cortactin | SRC8_HUMAN | 62 kDa | 7.6 | 17.5 ($\pm 0$) |
| 18 | Myosin light polypeptide 6 | MYL6_HUMAN | 17 kDa | 2.4 | 17.2 ($\pm 0.3$) |
| 19 | Nuclear pore complex protein Nup153 | NU153_HUMAN | 154 kDa | 21.9 | 17 ($\pm 0.5$) |
| 20 | Heterogeneous nuclear<br>ribonucleoprotein K | HNRPK_HUMAN | 51 kDa | 6.5 | 16.3 ($\pm 3$) |
| 21 | E3 SUMO-protein ligase RanBP2 | RBP2_HUMAN | 358 kDa | 55.9 | 15.4 ($\pm 3$) |
| 22 | Pre-mRNA-splicing factor SYF1 | SYF1_HUMAN | 100 kDa | 16.1 | 15.3 ($\pm 0$) |
| 23 | Stromal interaction molecule 1 | STIM1_HUMAN | 77 kDa | 12 | 14.7 ($\pm 0$) |
| 24 | Plakophilin-3 | PKP3_HUMAN | 87 kDa | 10.6 | 14.2 ($\pm 5.9$) |
| 25 | Beta/gamma crystallin domain-containing<br>protein 1 | CRBG1_HUMAN | 189 kDa | 20.5 | 14.2 ($\pm 7.4$) |
| 26 | Sickle tail protein homolog | SKT_HUMAN | 214 kDa | 26.9 | 14 ($\pm 0$) |
| 27 | Arf-GAP domain and FG repeat-containing protein 1 | AGFG1_HUMAN | 58 kDa | 12.7 | 13.3 ($\pm 1.4$) |
| 28 | Annexin A1 | ANXA1_HUMAN | 39 kDa | 5.1 | 12.4 ($\pm 4.2$) |
| 29 | Heterogeneous nuclear ribonucleoproteins A2/B1 | ROA2_HUMAN | 37 kDa | 6 | 12.2 ($\pm 0.4$) |
| 30 | Transmembrane protein 263 | TM263_HUMAN | 12 kDa | 2.4 | 11.8 ($\pm 1.1$) |
| 31 | Lymphoid-specific helicase | HELLS_HUMAN | 97 kDa | 9.9 | 11.1 ($\pm 1.1$) |
| 32 | Collagen alpha-1(XVII) chain | COHA1_HUMAN | 150 kDa | 18.4 | 10 ($\pm 0$) |
| 33 | Prelamin-A/C | LMNA_HUMAN | 74 kDa | 7.2 | 9.5 ($\pm 7.6$) |
| 34 | Plakophilin-1 | PKP1_HUMAN | 83 kDa | 5.5 | 9.5 ($\pm 0$) |
| 35 | Protein ELYS | ELYS_HUMAN | 253 kDa | 17.7 | 9.1 ($\pm 5.8$) |
| 36 | Rho GTPase-activating protein 21 | RHG21_HUMAN | 217 kDa | 15.1 | 8.9 ($\pm 1.9$) |
| 37 | Catenin alpha-1 | CTNA1_HUMAN | 100 kDa | 6.5 | 8.9 ($\pm 9.9$) |
| 38 | Protein FAM83H | FA83H_HUMAN | 127 kDa | 7.9 | 8.6 ($\pm 0$) |
| 39 | Delta (14)-sterol reductase LBR | LBR_HUMAN | 71 kDa | 9.3 | 7.9 ($\pm 0.9$) |
| 40 | Tubulin beta-4B chain | TBB4B_HUMAN | 50 kDa | 2.8 | 7.5 ($\pm 0$) |
| 41 | Heat shock cognate 71 kDa protein | HSP7C_HUMAN | 71 kDa | 5.7 | 7.4 ($\pm 0$) |
| 42 | Caveolin-1 | CAV1_HUMAN | 20 kDa | 1 | 7.1 ($\pm 0.8$) |
| 43 | RelA-associated inhibitor | IASPP_HUMAN | 89 kDa | 5.1 | 7 ($\pm 0.8$) |
| 44 | Epidermal growth factor receptor | EGFR_HUMAN | 134 kDa | 8.2 | 6.9 ($\pm 3.6$) |
| 45 | Annexin A2 | ANXA2_HUMAN | 39 kDa | 3.5 | 6.6 ($\pm 1.3$) |
| 46 | Microtubule-associated protein 4 | MAP4_HUMAN | 121 kDa | 5.5 | 6.4 ($\pm 0.1$) |
| 47 | Ladinin-1 | LAD1_HUMAN | 57 kDa | 2.3 | 6.1 ($\pm 1.2$) |
| 48 | Misshapen-like kinase 1 | MINK1_HUMAN | 150 kDa | 5.8 | 5.7 ($\pm 0$) |
| 49 | Elongation factor 1-alpha 1 | EF1A1_HUMAN | 50 kDa | 3.2 | 5.7 ($\pm 1.1$) |
| 50 | Heterogeneous nuclear<br>ribonucleoprotein M | HNRPM_HUMAN | 78 kDa | 4.1 | 5.7 ($\pm 7$) |

<sup>1</sup>Rank based on normalized percent coverage.
<sup>2</sup>Spectral counts from three biological replicates were normalized to endogenously biotinylated cellular proteins then averaged. Averages of the no-biotin control samples were subtracted from the plus-biotin experimental samples to determine the normalized spectral count.
<sup>3</sup>Average of the percent coverage of three biological replicates of the no-biotin control samples was subtracted from the average of the percent coverage of the plus-biotin experimental samples to determine the normalized percent coverage.
<sup>4</sup>Standard deviation (SD) of the three plus-biotin biological replicates.
<sup>5</sup>Peptides common to both PP1G\_HUMAN and PP1A\_HUMAN were identified as were peptides unique to each protein.

Gene ontogeny analysis of cellular proteins in proximity to tegument-delivered pUL21mT was performed using Metascape and is shown in Figure 8A. Prominent gene ontologies included cadherin binding, nuclear membrane, zonula adherens, and cell-substrate junction that may reflect the known roles of pUL21 in cell-to-cell spread of infection and nuclear egress [31, 35, 40, 41, 43–49]. Consistent with the subcellular localization of biotinylated proteins at 2 hpi (Figure 7), two of the top Metascape gene ontology categories were nuclear membrane and nuclear periphery. STRING analysis of protein-protein interaction networks of the cellular proteins identified 7 main interaction clusters including nuclear envelope breakdown, annexin, and zonula adherens (Figure 8B). In keeping with the established role of pUL21 as a PP1 adaptor protein [40, 62], it was interesting to identify the PTW/PP1 phosphatase complex in the STRING analysis. Of the 110 cellular proteins proximal to pUL21mT at 18 hpi, 35 were also in proximity to pUL21mT at 2 hpi (Supplementary Table 1). The top 10 unique proteins identified at 18 hpi and 2 hpi, based on normalized percent coverage, are listed in Supplementary Tables 2 and 3, respectively.

**Figure 8.**
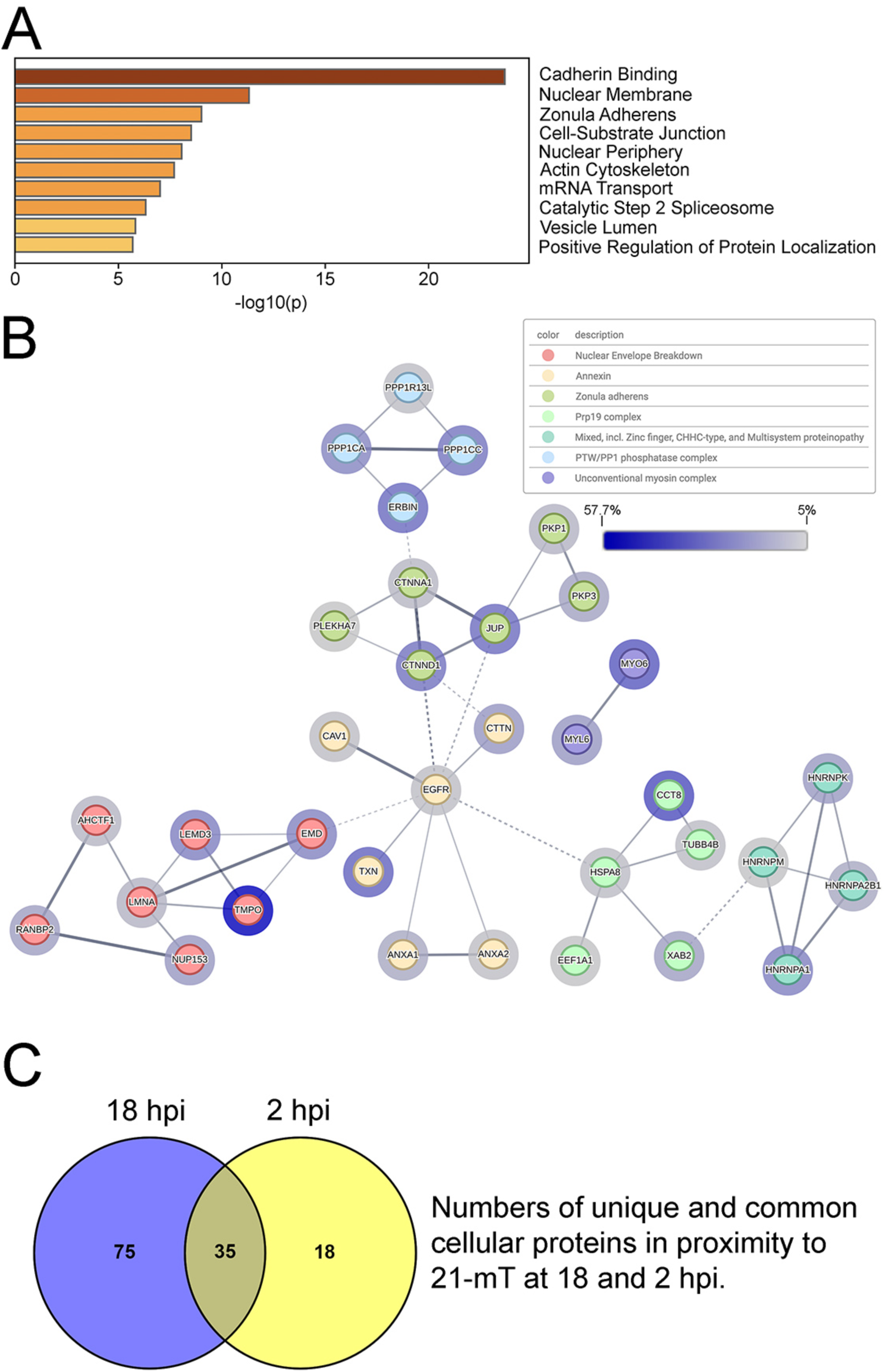
The identification of proteins in proximity to tegument-delivered pUL21mT immediately following viral entry. **(A)** Top 10 Metascape-generated gene ontologies of cellular proteins identified in proximity to tegument-delivered pUL21mT at two hours post-infection. **(B)** STRING network of cellular proteins in proximity to tegument-delivered pUL21mT upon viral entry. Identified proteins were analyzed, clustered, and colour-coded using the STRING database to visualize known protein-protein interactions. Edges connecting the nodes represent reported associations, with edge thickness corresponding to interaction confidence (thicker edges indicate higher confidence). Halos surrounding each node reflect percent coverage, with darker halos indicating higher percent coverage. Only proteins with a normalized protein coverage of ≥ 5% were included in this analysis, and disconnected nodes are not shown. **(C)** Venn diagram of the number of unique and common cellular proteins in proximity to pUL21mT at 18 and 2 hours post-infection (hpi).

Despite the success of leveraging BioID to identify proteins in proximity to pUL21mT during the early and late stages of infection, these putative interactions require further validation to determine their significance during HSV-2 infection. Erbin was uniquely identified as a proximal pUL21mT interactor at 2 hpi. Erbin is a multifunctional cellular protein that regulates inflammatory responses, cell signalling pathways such as the Ras/Raf/MEK/ERK mitogen-activated protein kinase (MAPK) pathway, and maintains epithelial cell-to-cell junction integrity [63–67]. Given Erbin’s broad activities and its ability to regulate cellular processes relevant to HSV-2 infection, we investigated the impact of pUL21 on Erbin expression. To do this, HaCaT cells were co-transfected with plasmids encoding HSV-2 pUL21 and myc-tagged Erbin [35, 68].

Erbin levels decreased when co-expressed with pUL21 in comparison with an empty vector control, suggesting that pUL21 mediates the degradation of Erbin in co-transfected cells (Figure 9). To explore the mechanism by which this occurs, transfected cells were treated with the proteasome inhibitor, MG-132 (Figure 9A), or the autophagy inhibitor, bafilomycin A (Figure 9B). Treatment with MG-132 or bafilomycin A failed to restore Erbin protein levels to those of the DMSO control. These data suggest that pUL21-mediated degradation of Erbin occurs independently of proteasomal and autophagy-mediated pathways, or alternatively, that redundancy between pathways allows Erbin to be degraded when one pathway is inhibited.

**Figure 9.**
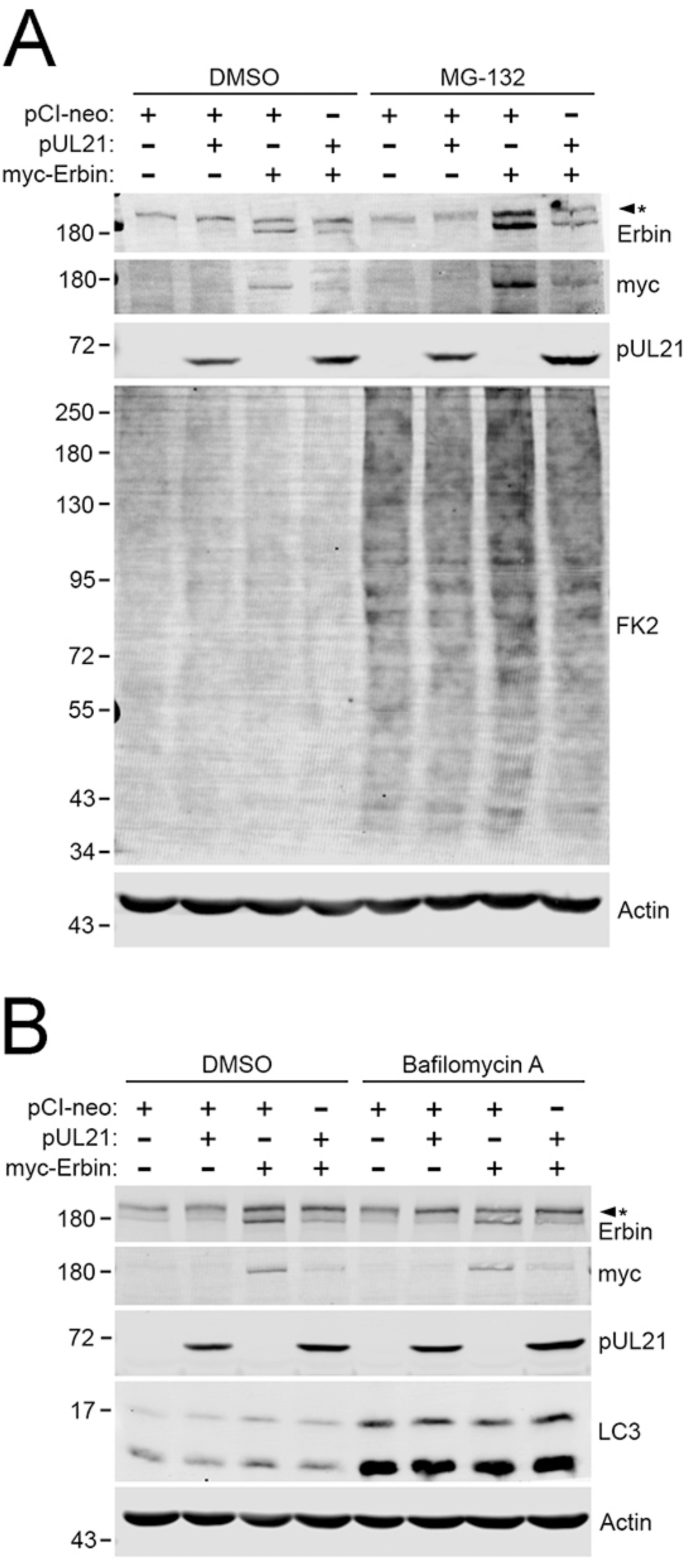
pUL21 mediates the degradation of Erbin. **(A)** pUL21-mediated degradation of Erbin is unaffected by proteasome inhibition. HaCaT cells were transfected with the indicated expression plasmids. At 18 hours post-transfection, cells were treated with DMSO or 10 µM MG-132 for 6 hours and analyzed by western blotting using antibodies against Erbin, myc-tag, pUL21, FK2 to identify ubiquitinated proteins, and actin. The asterisk indicates a background band reactive with the Erbin antisera. **(B)** pUL21-mediated degradation of Erbin is unaffected by inhibition of autophagy. HaCaT cells were transfected with the indicated expression plasmids. At 12 hours post-transfection, cells were treated with DMSO or 50 nM bafilomycin A for 12 hours, and analyzed by western blotting using antibodies against Erbin, myc-tag, pUL21, LC3, and actin. The asterisk indicates a background band reactive with the Erbin antisera. Migration positions of molecular weight markers (kDa) are shown on the left side of each western blot.

### Identification of proteins affinity-purified with pUL21 from infected cells

While BioID approaches are valuable for identifying molecules in proximity to a protein of interest in the context of living cells, a limitation of this technique is that one cannot distinguish between transient and stable interactions. To address this concern, we analyzed an HSV-2 strain that expresses pUL21 fused to mCherry (21-mCh) [42].

HaCaT cells were infected in triplicate with 21-mCh for 18 hours and pUL21mCh affinity-purified using RFP-TRAP beads. Proteins on beads were digested with trypsin and the resulting peptides were identified by LC-MS/MS. A total of 1545 cellular and viral proteins were identified in this analysis (supplemental data 3). These were further distilled as described above for the 18 hpi BioID experiments. Based on these criteria, 502 cellular and 62 viral proteins were further analyzed. Tables 4 and 5 list the top 50 cellular and viral proteins affinity-purified with pUL21mCh, ranked by averaged percent coverage.

**Table 4.** Top 50 Cellular Proteins Affinity-Purified With pUL21mCh at 18 hpi.

| <sup>1</sup>Rank | Protein Name | Gene Name | Molecular Weight | <sup>2</sup>Averaged<br>Spectral Count | <sup>3</sup>Averaged<br>Percent Coverage (±SD) |
| --- | --- | --- | --- | --- | --- |
| 1 | Serine/threonine-protein<br>phosphatase PP1-alpha<br>catalytic subunit | PP1A_HUMAN | 38 kDa | 94.7 | 79.3 (±7.2) |
| 2 | Tubulin beta-4B chain | TBB4B_HUMAN | 50 kDa | 92 | 76.8 (±5.3) |
| 3 | Serine/threonine-protein<br>phosphatase PP1-beta<br>catalytic subunit | PP1B_HUMAN | 37 kDa | 90.7 | 71.6 (±3.2) |
| 4 | Tubulin beta chain | TBB5_HUMAN | 50 kDa | 88 | 71.5 (±10.3) |
| 5 | Peptidyl-prolyl cis-trans<br>isomerase A | PPIA_HUMAN | 18 kDa | 18.7 | 69.3 (±4.5) |
| 6 | Annexin A2 OS=Homo<br>sapiens | ANXA2_HUMAN | 39 kDa | 49 | 68.4 (±0.3) |
| 7 | Actin, cytoplasmic 1 | ACTB_HUMAN | 42 kDa | 115 | 68.0 (±2.1) |
| 8 | Ras-related protein Rab-7a | RAB7A_HUMAN | 23 kDa | 15.3 | 66.4 (±4.0) |
| 9 | 14-3-3 protein sigma | 1433S_HUMAN | 28 kDa | 31.3 | 65.7 (±7.5) |
| 10 | Pyruvate kinase PKM | KPYM_HUMAN | 58 kDa | 55 | 65.3 (±1.4) |
| 11 | ATP synthase subunit beta,<br>mitochondrial | ATPB_HUMAN | 57 kDa | 50 | 65.1 (±5.8) |
| 12 | Triosephosphate isomerase | TPIS_HUMAN | 27 kDa | 16 | 64.1 (±6.4) |
| 13 | Voltage-dependent anion-<br>selective channel protein 1 | VDAC1_HUMAN | 31 kDa | 22 | 63.1 (±4.9) |
| 14 | Tubulin alpha-1B chain<br>OS=Homo sapiens | TBA1B_HUMAN | 50 kDa | 59 | 62.9 (±7.4) |
| 15 | Receptor of activated<br>protein C kinase 1 | RACK1_HUMAN | 35 kDa | 19 | 62.9 (±8.1) |
| 16 | Tubulin alpha-1C chain<br>OS=Homo sapiens | TBA1C_HUMAN | 50 kDa | 56 | 62.7 (±7.4) |
| 17 | Annexin A1 OS=Homo<br>sapiens | ANXA1_HUMAN | 39 kDa | 29 | 62.2 (±3.8) |
| 18 | Elongation factor 1-alpha 1<br>OS=Homo sapiens | EF1A1_HUMAN | 50 kDa | 72.3 | 61.2 (±2.0) |
| 19 | 14-3-3 protein epsilon | 1433E_HUMAN | 29 kDa | 26 | 60.9 (±4.0) |
| 20 | Serine/threonine-protein<br>phosphatase PP1-gamma<br>catalytic subunit | PP1G_HUMAN | 37 kDa | 85.3 | 60.7 (±5.9) |
| 21 | Prelamin-A/C | LMNA_HUMAN | 74 kDa | 72.3 | 58.4 (±2.1) |
| 22 | Protein disulfide-isomerase<br>A3 | PDIA3_HUMAN | 57 kDa | 45 | 57.2 (±2.8) |
| 23 | 3-hydroxyacyl-CoA<br>dehydrogenase type-2 | HCD2_HUMAN | 27 kDa | 13.7 | 57.0 (±3.8) |
| 24 | T-complex protein 1 subunit<br>theta | TCPQ_HUMAN | 60 kDa | 41.7 | 56.9 (±5.9) |
| 25 | T-complex protein 1 subunit<br>epsilon | TCPE_HUMAN | 60 kDa | 35.3 | 56.4 (±4.1) |
| 26 | Glyceraldehyde-3-phosphate dehydrogenase | G3P_HUMAN | 36 kDa | 32.3 | 55.6 (±2.8) |
| 27 | Tubulin alpha-4A chain | TBA4A_HUMAN | 50 kDa | 52.7 | 55.2 (±1.5) |
| 28 | T-complex protein 1 subunit<br>beta | TCPB_HUMAN | 57 kDa | 37 | 54.3 (±6.9) |
| 29 | Elongation factor Tu,<br>mitochondrial | EFTU_HUMAN | 50 kDa | 36 | 54.1 (±5.9) |
| 30 | Prohibitin OS=Homo<br>sapiens | PHB_HUMAN | 30 kDa | 16.3 | 53.8 (±13.0) |
| 31 | T-complex protein 1 subunit<br>gamma | TCPG_HUMAN | 61 kDa | 35 | 53.1 (±4.0) |
| 32 | Calreticulin OS=Homo sapiens | CALR_HUMAN | 48 kDa | 28 | 53.0 ($\pm 1.8$) |
| 33 | Malate dehydrogenase, mitochondrial | MDHM_HUMAN | 36 kDa | 28.3 | 52.0 ($\pm 3.4$) |
| 34 | Endoplasmic reticulum chaperone BiP | BIP_HUMAN | 72 kDa | 102.7 | 51.3 ($\pm 1.8$) |
| 35 | 14-3-3 protein zeta/delta | 1433Z_HUMAN | 28 kDa | 23 | 50.7 ($\pm 7.2$) |
| 36 | Fructose-bisphosphate aldolase A | ALDOA_HUMAN | 39 kDa | 16 | 50.6 ($\pm 12.3$) |
| 37 | Ras-related protein Rab-14 | RAB14_HUMAN | 24 kDa | 14 | 49.9 ($\pm 10.5$) |
| 38 | Profilin-1 | PROF1_HUMAN | 15 kDa | 7.7 | 49.8 ($\pm 7.7$) |
| 39 | Heterogeneous nuclear ribonucleoproteins A2/B1 | ROA2_HUMAN | 37 kDa | 26.7 | 49.5 ($\pm 1.8$) |
| 40 | T-complex protein 1 subunit delta | TCPD_HUMAN | 58 kDa | 31.3 | 49.1 ($\pm 3.9$) |
| 41 | ADP/ATP translocase 3 | ADT3_HUMAN | 33 kDa | 22.3 | 48.0 ($\pm 2.4$) |
| 42 | T-complex protein 1 subunit eta | TCPH_HUMAN | 59 kDa | 24 | 47.8 ($\pm 3.4$) |
| 43 | T-complex protein 1 subunit alpha | TCPA_HUMAN | 60 kDa | 35.7 | 46.7 ($\pm 5.2$) |
| 44 | Peroxiredoxin-1 OS=Homo sapiens | PRDX1_HUMAN | 22 kDa | 10.7 | 46.6 (±12.4) |
| 45 | Translationally-controlled tumor protein | TCTP_HUMAN | 20 kDa | 4 | 46.3 (±4.7) |
| 46 | Aldehyde dehydrogenase, dimeric NADP-preferring | AL3A1_HUMAN | 50 kDa | 36.3 | 46.3 (±5.2) |
| 47 | Myosin light polypeptide 6 | MYL6_HUMAN | 17 kDa | 7.3 | 45.5 (±8.4) |
| 48 | Cytoskeleton-associated protein 4 | CKAP4_HUMAN | 66 kDa | 37 | 45.2 (±3.4) |
| 49 | Y-box-binding protein 1 | YBOX1_HUMAN | 36 kDa | 11.3 | 45.1 (±3.6) |
| 50 | Protein disulfide-isomerase | PDIA1_HUMAN | 57 kDa | 26.3 | 44.6 (±11.7) |

<sup>1</sup>Rank based on the averaged percent coverage.
<sup>2</sup>Spectral counts from three biological replicates were averaged.
<sup>3</sup>Average of the percent coverage of three biological replicates.

**Table 5.** Top 50 Viral Proteins Affinity-Purified With pUL21mCh at 18 hpi.

| <sup>1</sup>Rank | Protein Name | Gene Name | Molecular Weight | <sup>2</sup>Averaged Spectral Count | <sup>3</sup>Averaged Percent Coverage (±SD) |
| --- | --- | --- | --- | --- | --- |
| 1 | Tegument protein UL21 | TEG4_HHV2H | 57 kDa | 1020.7 | 89.6 (±1.0) |
| 2 | Major DNA-binding protein | DNBI_HHV2H | 128 kDa | 807.3 | 80.7 (±0.4) |
| 3 | DNA polymerase processivity factor | PAP_HHV2H | 50 kDa | 141.7 | 77.8 (±4.7) |
| 4 | Triplex capsid protein 2 | TRX2_HHV2H | 34 kDa | 153.3 | 72.6 (±4.7) |
| 5 | Ribonucleoside-diphosphate reductase large subunit | RIR1_HHV23 | 125 kDa | 192 | 68.8 (±1.3) |
| 6 | Small capsomere-interacting protein | SCP_HHV2H | 12 kDa | 6.7 | 66.7 (±5.5) |
| 7 | Tegument protein UL51 | TEG7_HHV2H | 26 kDa | 19.3 | 63.5 (±7.1) |
| 8 | Major capsid protein | MCP_HHV2H | 149 kDa | 874.3 | 63.1 (±2.8) |
| 9 | Virion protein US10 | US10_HHV2H | 33 kDa | 33.7 | 62.7 (±6.1) |
| 10 | Capsid vertex component 1 | CVC1_HHV2H | 75 kDa | 65.7 | 60.6 (±1.9) |
| 11 | Tegument protein VP22 | VP22_HHV2H | 32 kDa | 43.7 | 60.4 (±1.2) |
| 12 | Tegument protein UL47 | TEG5_HHV2H | 74 kDa | 95.7 | 59.7 (±2.2) |
| 13 | Deoxyuridine 5'-triphosphate nucleotidohydrolase | DUT_HHV2H | 39 kDa | 19.3 | 57.7 (±6.4) |
| 14 | Triplex capsid protein 1 | TRX1_HHV2G | 50 kDa | 162.7 | 56.4 (±1.8) |
| 15 | Inner tegument protein | ITP_HHV2H | 119 kDa | 135.7 | 56.2 (±4.2) |
| 16 | Cytoplasmic envelopment protein 2 | CEP2_HHV2H | 40 kDa | 57 | 54.0 ($\pm 0.3$) |
| 17 | Capsid vertex component 2 | CVC2_HHV2H | 64 kDa | 71.7 | 53.9 ($\pm 0.2$) |
| 18 | Capsid scaffolding protein | SCAF_HHV2H | 67 kDa | 82.7 | 52.5 ($\pm 1.5$) |
| 19 | Ribonucleoside-diphosphate reductase small subunit | RIR2_HHV23 | 38 kDa | 26 | 52.4 ($\pm 4.9$) |
| 20 | Major viral transcription factor ICP4 homolog | ICP4_HHV2H | 135 kDa | 175 | 52.0 ($\pm 7.4$) |
| 21 | Protein US2 | US02_HHV2H | 33 kDa | 24.3 | 51.5 ($\pm 4.4$) |
| 22 | Serine/threonine-protein kinase US3 | US03_HHV2H | 53 kDa | 29 | 48.6 ($\pm 7.9$) |
| 23 | DNA polymerase catalytic subunit | DPOL_HHV21 | 137 kDa | 119 | 48.4 ($\pm 1.1$) |
| 24 | Envelope glycoprotein B | GB_HHV23 | 100 kDa | 101 | 47.8 ($\pm 1.8$) |
| 25 | Envelope glycoprotein C | GC_HHV23 | 52 kDa | 45 | 47.4 ($\pm 5.2$) |
| 26 | Nuclear egress protein 2 | NEC2_HHV2H | 30 kDa | 37.3 | 47.0 ($\pm 11.1$) |
| 27 | Uracil-DNA glycosylase | UNG_HHV2H | 28 kDa | 18.3 | 45.5 ($\pm 3.8$) |
| 28 | Large tegument protein deneddylase | LTP_HHV2H | 330 kDa | 160.3 | 45.2 ($\pm 4.7$) |
| 29 | Virion host shutoff protein | SHUT_HHV2G | 55 kDa | 24.3 | 45.2 ($\pm 1.4$) |
| 30 | Tegument protein VP16 | VP16_HHV23 | 55 kDa | 38.7 | 44.0 ($\pm 0.2$) |
| 31 | Envelope glycoprotein D | GD_HHV2H | 43 kDa | 59 | 43.7 ($\pm 0.7$) |
| 32 | mRNA export factor | ICP27_HHV2H | 55 kDa | 42.7 | 42.9 ($\pm 3.1$) |
| 33 | Portal protein | PORTL_HHV2H | 75 kDa | 37.7 | 42.7 ($\pm 2.0$) |
| 34 | Envelope glycoprotein H | GH_HHV2H | 90 kDa | 49.3 | 42.3 ($\pm 1.8$) |
| 35 | Envelope glycoprotein G | GG_HHV2B | 72 kDa | 96 | 42.2 ($\pm 0.6$) |
| 36 | Alkaline nuclease | AN_HHV2 | 66 kDa | 42.7 | 42.1 ($\pm 2.7$) |
| 37 | Tripartite terminase subunit 3 | TRM3_HHV2H | 81 kDa | 28.7 | 40.8 ($\pm 6.2$) |
| 38 | Thymidine kinase | KITH_HHV23 | 40 kDa | 23.7 | 39.8 ($\pm 2.1$) |
| 39 | Replication origin-binding protein | OBP_HHV2H | 96 kDa | 49.7 | 39.5 ($\pm 3.3$) |
| 40 | Nuclear egress protein 1 | NEC1_HHV2H | 34 kDa | 20.7 | 39.0 ($\pm 2.6$) |
| 41 | Tegument protein UL14 | TEG3_HHV2H | 24 kDa | 5 | 38.5 ($\pm 4.3$) |
| 42 | Tegument protein UL46 | TEG1_HHV2H | 78 kDa | 37.7 | 37.7 ($\pm 8.8$) |
| 43 | Tegument protein UL55 | TEG6_HHV2H | 20 kDa | 8 | 37.1 ($\pm 5.5$) |
| 44 | Envelope glycoprotein I | GI_HHV23 | 40 kDa | 26.3 | 36.3 ($\pm 0.0$) |
| 45 | DNA primase | PRIM_HHV2H | 115 kDa | 34.7 | 36.1 ($\pm 2.2$) |
| 46 | Envelope glycoprotein M | GM_HHV2H | 50 kDa | 17.7 | 32.4 ($\pm 5.3$) |
| 47 | Cytoplasmic envelopment protein 1 | CEP1_HHV2H | 33 kDa | 14.7 | 31.8 ($\pm 2.0$) |
| 48 | Probable RNA-binding protein | RNB_HHV2H | 16 kDa | 12.3 | 30.0 ($\pm 6.0$) |
| 49 | E3 ubiquitin ligase ICP22 | ICP22_HHV2H | 45 kDa | 23.3 | 28.0 ($\pm 4.5$) |
| 50 | DNA replication helicase | HELI_HHV2H | 99 kDa | 20.3 | 24.4 ($\pm 1.3$) |

<sup>1</sup>Rank based on the averaged percent coverage.
<sup>2</sup>Spectral counts from three biological replicates were averaged.
<sup>3</sup>Average of the percent coverage of three biological replicates.

Gene ontogeny analysis of cellular proteins that were affinity-purified with pUL21mCh was performed using Metascape and is shown in Figure 10A. Many proteins involved in cadherin binding, mRNA metabolism, cell-substrate junctions, and energy metabolism were identified. STRING 12.0 analysis of protein-protein interaction networks of the cellular proteins identified 12 main interaction clusters including folding of cytoskeletal components, the spliceosome, energy metabolism, nuclear pore localization, and nuclear membrane-associated proteins (Figure 10B).

**Figure 10.**
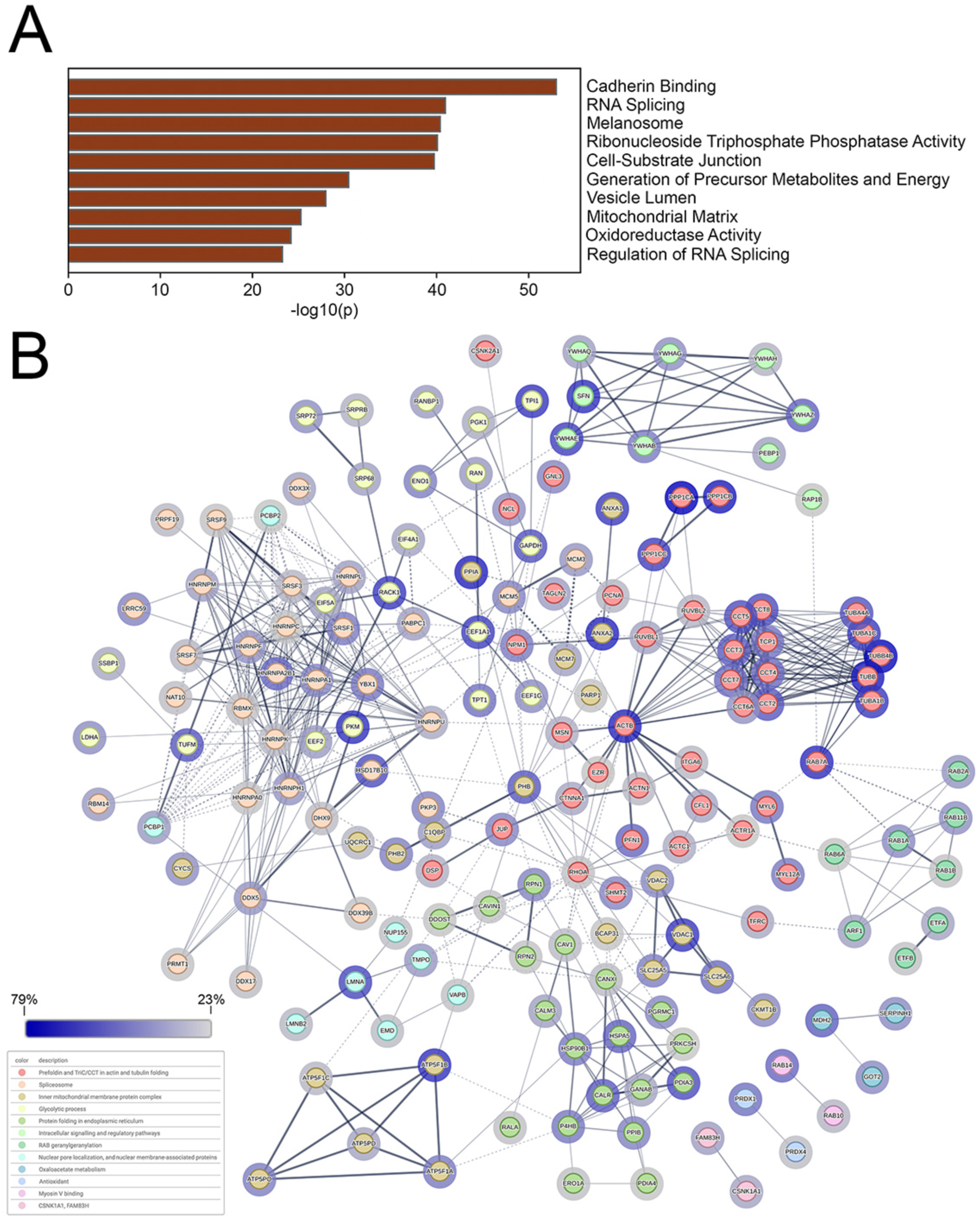
The identification of proteins affinity-purified with pUL21mCh in infected HaCaT cells. **(A)** Top 10 Metascape-generated gene ontologies of cellular proteins affinity-purified with pUL21mCh at 18 hours post-infection. **(B)** STRING network of cellular proteins associated with pUL21mCh at 18 hours post-infection. The top 200 cellular proteins, ranked by average percent coverage, were analyzed, clustered, and colour-coded using the STRING database to visualize known protein-protein interactions. Edges connecting the nodes represent reported associations, with edge thickness corresponding to interaction confidence (thicker edges indicate higher confidence). Halos surrounding each node reflect percent coverage, with darker halos indicating higher percent coverage and disconnected nodes are not shown.

Comparison of our 18 hpi BioID results with those obtained by affinity purification of pUL21mCh at 18 hpi identified 46 common cellular proteins (Supplementary Table 4, Supplementary Figure 1A) and 14 common viral proteins (Supplementary Table 5, Supplementary Figure 1C). Comparison of the 2 hpi BioID results with the affinity purification data identified 28 common cellular proteins (Supplementary Table 6, Supplementary Figure 1B). STRING analysis of cellular proteins identified by both methods at 18 hpi yielded seven conserved clusters including adherens junctions, the spliceosome, and the nuclear envelope (Supplementary Figure 2).

## Discussion

Utilizing multiple approaches, we have performed a comprehensive analysis of proteins in association with the multifunctional tegument protein, pUL21, at early and late stages of HSV-2 infection. To identify proteins in proximity to pUL21, we turned to BioID, whereby a “bait” protein of interest (e.g. pUL21) is fused to a promiscuous biotin ligase, such as the third-generation enzyme mT, that is much more efficient and smaller than the BirA* and BioID2 enzymes [53]. When cells expressing the mT-fusion are incubated with biotin, mT produces a short-lived reactive biotin species that becomes covalently linked to proximal lysine residues within an approximately 10nm radius [51]. Cells are lysed and biotinylated proteins are captured on streptavidin-conjugated beads that, due to the strength of the biotin-avidin interaction, can withstand washing with buffers containing salts and detergents at concentrations that would interfere with both specific and non-specific protein-protein interactions. Thus, the BioID approach has the advantage of a low non-specific background, improving confidence in the identified proteins. Additionally, BioID can be used to identify weak or transient interactions between bait and prey proteins that would not be identified by co-affinity purification methodologies. Importantly, BioID offers the advantage of probing proximal protein interactions in the context of living cells. As BioID relies on protein proximity rather than protein-protein interactions, direct or indirect contacts between bait and labelled prey proteins cannot be implied. Furthermore, as biotin labelling depends on the presence of proximal lysine residues, proteins with low lysine content will be underrepresented, or absent, in BioID analyses. Affinity purification enables the identification of proteins in direct or indirect contact with the tagged bait protein, but suffers from the identification of non-specific interactions, artifactual interactions initiated after cell lysis, the loss of weak interactions, and the failure to capture interactions in the context of intact cells.

Thus, we, as have others [52, 69], chose to take advantage of the complementary nature of BioID and affinity purification to gain a broader view of proteins associating with pUL21, with the rationale that pUL21-associated proteins identified by both methodologies would represent stronger candidates for *bona fide* pUL21 interaction partners compared to those identified using a single approach.

A total of 46 cellular proteins were identified in both BioID and affinity purification analysis at 18 hpi (Figure 11A, Table 5). Of these, 33 were found within 7 interaction clusters, including phosphatase, adherens junctions, nuclear envelope, and spliceosome. The protein phosphatase catalytic subunits PP1-alpha and PP1-gamma were top hits identified in proximity to HSV-2 pUL21mT at 2 hpi and 18 hpi (Supplementary Table 1) and were in complex with HSV-2 pUL21mCh at 18 hpi (Table 4). Given the extensive overlap in peptide sequences among the alpha, beta and gamma PP1 isoforms, we cannot exclude the possibility that PP1-beta is also in association with HSV-2 pUL21. We identified multiple PP1 peptides that were common to all three isoforms and a few that were unique to the alpha and gamma isoforms.

Based on the findings of Graham and colleagues that characterized the interaction of HSV-1 pUL21 with PP1 in considerable detail [40, 62, 70], this result was not unexpected and serves as an important validation of our approach. pUL21 is a PP1 adapter protein that directs PP1 phosphatase activity towards both cellular and viral substrates [40, 62]. Binding of PP1 by pUL21 appears to be critical for HSV-1 replication and spread, as well as the dephosphorylation of substrates of the viral serine/threonine kinase pUs3 [40].

Analysis of numerous pUL21 deletion mutants in HSV-1 and HSV-2 strains has shown that pUL21 functions in cell-to-cell spread of virus infection [35, 40, 43–49]. How pUL21 promotes cell-to-cell spread of infection is not well understood; however, in association with pUL16 and pUL11, pUL21 is assembled into a complex on the cytoplasmic tail of glycoprotein E (gE) that is required for the efficient delivery of gE to cell surfaces [48]. gE is well known to traffic to adherens junctions [71] and recruit nascent enveloped virions to cell junctions to promote cell-to-cell spread of infection [72]. The delivery of pUL21 to these sites provides opportunities for pUL21 to interact with components of cell junctions and modulate their activities; perhaps through PP1-mediated dephosphorylation of junctional components, which is well known to regulate intercellular junction activities [73]. Alternatively, it may be that the pUL21 function in cell-to-cell spread is limited to facilitating appropriate trafficking of gE to sites of cell-to-cell spread; however, findings from the Wills group suggest a more complex role for pUL21 in cell-to-cell spread, demonstrating that pUL21 is required for syncytia formation mediated by syncytial mutants in glycoprotein B (gB), but not in other *syn* loci (i.e. UL53 (gK), UL20 and UL24) [48, 49].

Interestingly, the INM protein lamina-associated polypeptide 2, isoforms beta/gamma (LAP2γ/ψ) was the top cellular protein in proximity to pUL21mT at both 18 and 2 hpi (Tables 1 and 3) and was also readily detected in complex with pUL21mCh at 18 hpi (35.6% coverage, 14.3 average spectral counts, supplementary data 3). Other INM proteins identified by BioID included the cellular proteins Man1, delta (14)-sterol reductase LBR, SUN domain containing protein 1, and emerin, as well as the viral NEC components pUL31 and pUL34. The nuclear lamina component, prelamin A/C, which abuts the INM, was also in proximity to pUL21. We have previously reported that HSV-2 pUL21 localizes to the nuclear envelope, where it regulates nuclear egress at the INM and associates with NPCs on the cytoplasmic face of the nuclear envelope to prevent the binding of nascent cytoplasmic nucleocapsids during the late stages of infection [38, 42]. However, the mechanism by which pUL21 is recruited to the INM is poorly understood. pUL21 localizes to the INM in the absence of other viral proteins (Figure 5) and lacks any canonical sequences that would facilitate its association with membranes suggesting that pUL21 associates with cellular proteins at the INM. Experiments investigating the biophysical properties of INM-associated pUL21 suggest that pUL21 has the properties of an INM protein rather than a component of the nuclear lamina (Figure 5). Thus, we hypothesize that interactions between pUL21 and one, or more, of the cellular INM proteins we identified in proximity to pUL21 mediate pUL21 recruitment to the INM. The NPC component, RanBP2 (also known as Nup358), that forms the filaments on the cytoplasmic face of NPCs, was identified in proximity to pUL21 at 2 hpi and 18 hpi. As RanBP2 has been implicated in recruiting incoming capsids to NPCs following virus entry [74], it is a strong candidate for mediating pUL21 inhibition of nascent capsid recruitment to NPCs late in infection.

Unexpectedly, many cellular components involved in mRNA splicing and metabolism were found to be associated with pUL21 late in infection. We are unaware of any reported pUL21 functions that might explain these associations. The C-terminus of HSV-1 pUL21 has been suggested to have RNA-binding activity, which may facilitate these interactions [75]; however, it should be noted that a more recent study failed to detect any pUL21 RNA-binding activity [62]. It will be of interest to explore the impact of pUL21 expression on mRNA splicing and the consequences that this may have on cellular gene expression in virally infected cells.

Ma and colleagues have reported that pUL21 from pseudorabies virus (PRV) and HSV-1 facilitates the selective autophagy of cyclic GMP-AMP synthase (cGAS), a critical cytoplasmic DNA sensor that, once activated, triggers the production of interferon and subsequent inhibition of virus replication [50]. Mechanistically, pUL21-mediated degradation of cGAS is facilitated by the N-terminal domain of pUL21 that recruits cGAS, the ubiquitin ligase, UBE3C, that ubiquitinates cGAS, and the selective autophagy cargo receptor, TOLLIP, that directs the complex to a phagophore membrane, leading to its degradation [50]. We did not find cGAS, UBE3C, or TOLLIP associated with HSV-2 pUL21 in our experiments. It may be that the efficient degradation of these molecules in infected cells precluded their identification in our studies. The inclusion of an autophagy inhibitor in future analyses may enable the identification of this complex and perhaps additional targets of pUL21-mediated selective autophagy. Whereas Benedyk and colleagues identified the sphingomyelin transport protein, ceramide transfer protein (CERT), as an HSV-1 pUL21-interacting molecule in HEK 293T cells, it was not identified in our studies using HaCaT cells [40, 62]. The sequences of HSV-1 pUL21 involved in CERT binding are conserved in HSV-2, and HSV-1 pUL21 regulation of CERT in HaCaT cells has been documented [62]; therefore, HSV species-specific differences or differences between cell lines are not likely responsible for this discrepancy.

In addition to the NEC components pUL31 and pUL34, a number of other viral proteins were commonly associated with pUL21 in both BioID and affinity purification experiments (Supplementary Table 4). These included several tegument proteins including pUL36, pUL37, pUL47, pUL49, and pUL51, as well as the capsid component pUL38. It will be of interest to determine if any of these proteins are involved in the recruitment of pUL21 to capsids. It was curious that the well-known pUL21-binding protein, pUL16, was not identified in our BioID experiments despite providing a robust signal in affinity purification experiments. We hypothesize that this discrepancy is due to the relative paucity of lysine residues incorporated into pUL16 (three lysine residues out of 372 amino acids) that are required for protein biotinylation.

Remarkably, using BioID, we were able to identify cellular proteins in proximity to tegument-delivered pUL21 prior to *de novo* viral protein synthesis. The HSV tegument is a complex subvirion compartment containing roughly 20 viral proteins and about 50 cellular proteins [1]. A widely held view is that the tegument proteins delivered to the host cell cytoplasm upon infection serve to alter the cellular landscape, making it more conducive to virus replication. For example, tegument-mediated delivery of the viral ribonuclease, vhs, degrades cellular mRNAs undergoing translation and thereby initiates the shutoff of host cell protein synthesis as a prelude to the robust expression of virus-encoded proteins [14, 76]. Following viral envelope fusion with a cellular membrane, the capsid-associated tegument protein, pUL36, associates with the cellular microtubule motor dynein-dynactin that is required for the initial stages of capsid transport towards the nucleus where the incoming viral genome is ultimately deposited (reviewed in [77]). Additionally, tegument-delivered VP16 interacts with host cell proteins, host cell factor-1 and Oct-1, forming a complex that is delivered to the nucleus where it stimulates the transcription of immediate early viral genes from incoming viral genomes [17, 18]. Aside from these three well-characterized examples, the activities of most tegument proteins immediately following their delivery to the cell cytoplasm are poorly understood. Studies from a number of laboratories have suggested that the majority of tegument proteins rapidly disperse upon virion envelope fusion with a cellular membrane, whereas a few, such as pUL36 and pUL37, which are critical for capsid delivery to the nucleus, remain capsid-associated [2–12]. Our findings support the idea that pUL21 also rapidly disperses upon viral entry as no virus-encoded proteins meeting our filtering criteria were identified in proximity to tegument-delivered pUL21, other than pUL21 itself. Furthermore, pUL21mT-labelled proteins localized predominantly at the nuclear rim and throughout the nucleus at early times post-infection, suggesting that incoming pUL21mT disseminated widely throughout the infected cell (Figure 7).

Amongst the cellular proteins uniquely identified in proximity to pUL21 at 2 hpi (Table 6), the ERBB2-interacting protein, Erbin, which was identified with an average of 37.8 spectral counts covering roughly 30% of the 158 kDa protein, caught our attention. Erbin is a leucine-rich repeat-containing (LRR) protein that functions as a signalling scaffold [78]. LRR-containing proteins have diverse functions but are commonly found in molecules that function in innate and adaptive immune responses [79]. Erbin, a plasma membrane-associated protein that can localize to adherens junctions, inhibits the Ras/Raf/MEK/ERK MAP kinase pathway by binding to activated Ras, preventing its interaction with Raf and subsequent Raf/MEK/ERK activation [64, 80]. Prohibitin 1 (PHB1), a plasma membrane-associated scaffolding protein that binds to Raf and facilitates its activation and the downstream activation of MEK and ERK, binds to gE in HSV-1 infected cells and is required for the gE-dependent/ERK-dependent cell-to-cell spread of HSV-1 at cell junctions [81]. While PHB1 promotes ERK activation at cell junctions, Erbin directly opposes PHB1 activity by preventing the activation of ERK at cell junctions. In addition to cell-to-cell spread, an emerging role for ERK activity at gap junctions has been recently recognized. The Favoreel group determined that PRV infection triggers ERK-mediated phosphorylation of the gap junction component connexin 43, thereby closing gap junctions. Gap junction closure was suggested to prevent the intercellular diffusion of small biomolecules such as cyclic GMP-AMP that, via STING, function in innate immune responses to viral infection and might be expected to hinder virus transmission between cells [82]. Thus, there is a growing body of evidence suggesting the existence of proviral ERK activities at cell junctions. It follows that repression of ERK activation at cell junctions, mediated by Erbin, for example, might be viewed as antiviral. Our observations that pUL21 expression leads to the degradation/destabilization of Erbin (Figure 9) suggest that pUL21 could be antagonizing an Erbin-mediated antiviral response. As pUL21-mediated reductions in Erbin levels appear to be independent of proteasomal or autophagy mechanisms alone, it will be of interest to determine whether pUL21/PP1 influences the phosphorylation status of Erbin, thereby affecting Erbin stability or activity in virus-infected cells.

In summary, we have identified cellular and viral proteins associated with HSV-2 pUL21 at early and late stages of virus infection of human keratinocytes. Some of the proteins identified, such as PP1 isoforms were expected, and many, while not expected, were not unanticipated based on the known activities of pUL21 in nuclear egress and cell-to-cell spread of infection. Further investigation of this latter group of proteins promises to advance our understanding of biological processes that are critical for the replication of all herpesviruses. Some of the proteins identified were completely unexpected and the robustness of their common ontological associations suggests that pUL21 is very likely involved in heretofore unappreciated processes, such as the regulation of mRNA splicing. Finally, our demonstration that mT fusions to tegument proteins enables the identification of proteins associating with these virion components immediately after infection provides a powerful new approach for understanding the consequences of tegument delivery to the newly infected cell.

## Materials and Methods

### Cells and viruses

Human keratinocytes (HaCaT), HaCaT cells stably expressing pUL21 (HaCaT21) [46], and HeLa cells were maintained in Dulbecco’s modified Eagle medium (DMEM) supplemented with 10% fetal bovine serum (FBS) in a 5% CO_2_ environment. HSV-2 strain 186 lacking pUL21 (ΔUL21) and HSV-2 strain 186 carrying pUL21 fused to mCherry (21-mCh) have been described previously [35, 44]. Times post-infection, reported as hours post-infection (hpi) refers to the time elapsed following a one-hour inoculation period.

HSV-2 strain 186 expressing pUL21 fused to miniTurbo to its carboxy terminus (21-mT) was constructed by two-step Red-mediated mutagenesis [55] using a bacterial artificial chromosome (BAC) clone pYEbac373 [35] in *Escherichia coli* GS1783. Primer 5’ GCT TAC CGT TTG CCT GGC TCG CGC CCA GCA CGG CCA GTC TGT GGC TAG CAT CCC GCT GCT GAA CGC 3’ and 5’ ATC CGT GGG TTA GAA AAC GAC TGC ACT TTA TTG GGA TAT CTC AGT CGG CCC TGC TGA ATT CC 3’ were used to amplify a PCR product from pEP-Kan-S2 encoding 3x HAminiTurbo-in and used to add the miniTurbo coding sequence directly to the 3’ end of UL21. Restriction fragment length polymorphism analysis was used to confirm the integrity of each BAC clone in comparison to the WT BAC by digestion with *Eco*RI. Additionally, a PCR fragment spanning the mutated region of interest was amplified and sequenced to confirm the appropriate protein fusion. Infectious virus was reconstituted from BAC DNA as described previously [35].

### Plasmids and transfections

To construct pEP-Kan-S2 3xHAminiTurbo-in, HAminiTurbo163 (aa 1-163) was PCR amplified using primers 5’ GATCGGATCCATGTACCCGTATGATGTTCCGG 3’ and 5’ GATCGGATCCTGTTATTCCGGCCAGCTCCACC 3’, and HAminiTurbo147stop (aa 147-stop codon) was PCR amplified using primers 5’ GATCGCGGCCGCCTGTATCTGCAGGATAGAAAGC 3’ and 5’ GATCGCGGCCGCTCAGTCGGCCCTGCTGAATTCCTTTTCG 3’. 3xHA-miniTurbo-NLS_pCDNA3 (Addgene, Watertown, MA) was used as the template for both. The two fragments were sequentially cloned by first cloning HAminiTurbo163 digested with BamHI into similarly digested pEP-Kan-S2, a kind gift of Klaus Osterrieder (Freie Universität Berlin). The resulting plasmid was used to clone HAminiTurbo147stop digested with Not I into a similarly digested plasmid. All plasmids were sequenced to ensure that no spurious mutations were introduced during their construction. The construction of plasmids encoding HSV-2 pUL21 and HSV-2 pUL21-EGFP was described previously [35]. The construction of EGFP-UL31 and FLAG-UL34 was described previously [83]. Other plasmids used in this study were pFLAG-CMV-2 (Sigma Aldrich, St. Louis, MO), pCI-neo (Promega, Madison, WI), pClneoMyc human ERBIN [68] (Addgene plasmid # 40214 a gift from Yutaka Hata) and pGFP-LB1 [84], a gift from Dr. RD Goldman, Northwestern University). Plasmids were transfected into HeLa and HaCaT cells using X-tremeGene HP DNA transfection reagent (Roche, Laval, QC) following the manufacturer’s protocols.

### Immunological reagents and chemicals

Mouse monoclonal antibody against ICP27 (Virusys, Taneytown, MD) was used for western blotting and indirect immunofluorescence microscopy at a dilution of 1:500; mouse monoclonal antibody against γ-actin (Sigma) was used for western blotting at a dilution of 1:2,000; mouse monoclonal antibody against HSV ICP5 (VP5) (Virusys) was used for western blotting at a dilution of 1:500; mouse monoclonal antibody against BirA* (Novus Biologicals, Centennial, CO) was used for western blotting at a dilution of 1:500; rat polyclonal antiserum against HSV-2 pUL21 [35] was used for western blotting at a dilution of 1:600; rabbit polyclonal antibody against Erbin (Novus Biologicals) was used for western blotting at a dilution of 1:500; mouse monoclonal antibody against myc-tag (Roche) was used for western blotting at a dilution of 1:100; mouse monoclonal antibody against mono- and polyubiquitinylated conjugates (FK2; Enzo Life Sciences, Farmingdale, NY) was used for western blotting at a dilution of 1:1,000; rabbit polyclonal antibody against LC3 (Proteintech, Rosemont, IL) was used for western blotting at a dilution of 1:1,000; horseradish peroxidase (HRP)-conjugated streptavidin (SA-HRP; Invitrogen, Waltham, MA) was used for western blotting at a dilution of 1:100,000; HRP-conjugated goat anti-mouse (Jackson ImmunoResearch, West Grove, PA) and HRP-conjugated goat-anti rat (Sigma) were used for western blotting at a dilution of 1:30,000 and 1:80,000, respectively; IRDye 680RD Streptavidin (LI-COR Biosciences, Lincoln, NE) was used for western blotting at a dilution of 1:1,000; IRDye 680RD and 800CW anti-mouse secondary antibody, IRDye 680RD and 800CW anti-rat secondary antibody, and IRDye 680RD and 800CW anti-rabbit secondary antibody (Li-COR Biosciences) were used for western blotting at a dilution of 1:15,000; rat polyclonal antiserum against HSV-2 pUs3 [85] was used for indirect immunofluorescence microscopy at a dilution of 1:500; chicken polyclonal antiserum against HSV-2 pUL34 [41] was used for indirect immunofluorescence microscopy at a dilution of 1:500; Alexa Fluor 488-conjugated streptavidin (SA-488; Invitrogen) was used for indirect immunofluorescence microscopy at a dilution of 1:3,000; Alexa Fluor 568-conjugated donkey anti-chicken and Alexa Fluor 568-conjugated donkey anti-mouse (Molecular Probes, Eugene, OR) were used for indirect immunofluorescence microscopy at a dilution of 1:500. Biotin (Bio Basic, Markham, ON), Cycloheximide (CHx), MG-132, and bafilomycin A (Sigma) were used at 50 µM, 50 µg/mL, 10 µM, and 50 nM, respectively.

### Preparation of whole cell extracts for western blot analysis

To prepare whole cell extracts for western blot analysis, 4×10^6^ HaCaT cells were infected at a multiplicity of infection (MOI) of 1 or transfected, washed with cold phosphate-buffered saline (PBS), and scraped into 50-100 µL of PBS containing protease and phosphatase inhibitors (Roche), and transferred to microcentrifuge tubes containing 50 µL of 3X SDS-PAGE loading buffer. Transfected samples were treated with 250 units of benzonase nuclease (Santa Cruz Biotechnology, Dallas, TX) for 20 minutes at room temperature prior to the addition of 3X SDS-PAGE loading buffer. Cell lysates were passed through a 28 ½-gauge needle to reduce their viscosity, boiled for 10 minutes, and briefly centrifuged. For western blot analysis, 10-15 µL of cell lysates were electrophoresed through regular SDS-PAGE gels and separated proteins were transferred to polyvinylidene difluoride (PVDF) membranes (Millipore, Billerica, MA) and probed with appropriate dilutions of primary and secondary antibodies. Membranes were treated with Pierce ECL western blotting substrate (Thermo Scientific, Rockford, IL) and imaged using an Azure c300 Gel Imaging System. Alternatively, primary antibodies were detected with an appropriate IRDye-labelled secondary antibody and imaged using an Odyssey DLx Infrared Imaging System (LI-COR Biosciences).

### Virus replication analysis

4×10^6^ HaCaT cells were infected in triplicate at a multiplicity of infection (MOI) of 0.1 for endpoint titration analysis. After a one-hour inoculation period, cells were treated with low pH citrate buffer (40 mM Na citrate, 10 mM KCl, 0.8% NaCl) for 3 minutes at 37°C to inactivate extracellular virions, washed once with medium, and incubated at 37°C in complete medium, supplemented with biotin-depleted FBS with or without 50 µM of exogenous biotin. At 48 hpi, medium and cells were harvested, subjected to three freeze/thaw cycles, briefly sonicated in a chilled cup-horn sonicator, and centrifuged at 3000 rpm at 4°C to remove cellular debris. Titers were determined by plaque assay on HaCaT21 cells.

### Analysis of cell-to-cell spread of infection

Monolayers of HaCaT cells on 35 mm glass-bottom dishes (MatTek, Ashland, MA) were infected at various 10-fold dilutions required to yield well-isolated plaques and overlaid with complete medium containing 1% carboxymethylcellulose with or without biotin-depleted FBS. At 48 hpi, cells were fixed and permeabilized with ice-cold 100% methanol, and stained for the HSV viral kinase pUs3 as described previously [85]. Images of plaques (n=55 for each virus strain) were taken using a Nikon TE200 inverted epifluorescence microscope using a 10X objective and a cooled CCD camera. Plaque size was quantified by measuring plaque area using Image-Pro 6.3 software (Media Cybernetics, Bethesda, MD). Single infected cells were not included in this analysis.

### Preliminary analysis of pUL21mT biotinylation profile by western blotting

4×10^6^ HaCaT cells were infected at an MOI of 1. Following a one-hour inoculation period, cells were incubated at 37°C in complete medium, supplemented with biotin-depleted FBS. At 6, 10, or 18 hpi, cells were treated with 50 µM exogenous biotin for one hour and harvested for western blot analysis.

### Affinity purification of proteins biotinylated by pUL21mT

2.4×10^7^ HaCaT cells were infected at an MOI of 1. Following a one-hour inoculation period, cells were incubated at 37°C in complete medium, supplemented with biotin-depleted FBS and treated with 50 µM exogenous biotin for one hour at 18 hpi, or treated with 50 µg/mL CHx and 50 µM exogenous biotin between 0-2 or 2-4 hpi. Cells were then washed with ice-cold PBS, harvested in 1 mL ice-cold PBS, and centrifuged at 500 xg at 4°C for 5 minutes. Biotinylated proteins were immobilized on streptavidin-Sepharose beads (GE Healthcare, Chicago, IL) or Dynabeads MyOne Streptavidin C1 beads (Invitrogen) according to the manufacturer’s instructions, with modifications based on a previously published protocol [86]. Following incubation, bead-bound proteins were pelleted by centrifugation (Bound; B), and post-pull-down supernatants (S) were collected. A portion of each sample was retained prior to the addition of streptavidin-Sepharose beads (Input; I). Biotinylated proteins were visualized by western blot using SA-HRP or IRDye 680RD streptavidin.

### Indirect immunofluorescence microscopy

Infected and transfected cells were fixed at the indicated times post-infection or post-transfection with 4% paraformaldehyde in PBS for 15 minutes at room temperature. Cells were washed three times with PBS and permeabilized using 1% Triton X-100 in PBS for 10-30 minutes at room temperature. Following permeabilization, cells were washed three times with PBS/1% bovine serum albumin (BSA; PBS/1% BSA), incubated in primary antibody diluted in PBS/1%BSA for one hour at room temperature, washed three times with PBS/1% BSA, and incubated with appropriate Alexa Fluor conjugated secondary antibodies diluted in PBS/1% BSA for 0.5-1 hour at room temperature. After secondary antibody incubation, cells were washed three times with PBS/1% BSA, incubated with 0.5 µg/mL of Hoechst 33342 (Sigma) in PBS for 10 minutes at room temperature, and washed three times with PBS/1% BSA and stored at 4°C in PBS/1% BSA. Samples were imaged through a 60X (1.42 NA) oil immersion objective using an Olympus FV1000 confocal laser scanning microscope and FV10 ASW 4.01 software.

### Sample preparation for BioID mass spectrometry

6×10^7^ HaCaT cells were infected in triplicate at an MOI of 3. Following a one-hour inoculation period, cells were incubated at 37°C in complete medium, supplemented with biotin-depleted FBS and treated with 50 µM exogenous biotin for one hour at 18 hpi, or treated with 50 µg/mL CHx and 50 µM exogenous biotin between 0-2 hpi. Cells were then washed with ice-cold PBS, harvested in 1 mL ice-cold PBS, and centrifuged at 500 xg at 4°C for 5 minutes. Biotinylated proteins were immobilized on Dynabeads MyOne Streptavidin C1 beads (Invitrogen) according to the manufacturer’s instructions, with modifications based on a previously published protocol [86]. Bead-bound proteins were digested with trypsin, and orbitrap/iontrap tandem mass spectrometry was performed by the Southern Alberta Mass Spectrometry Facility at the University of Calgary. Tandem mass spectra data were searched against a human protein database containing WT HSV proteins.

### Sample preparation for proteins associated with pUL21mCh mass spectrometry

Lysates from HaCaT cells grown on two 150 mm dishes and infected with HSV-2 strain 186 viruses carrying pUL21-mCherry fusion proteins at an MOI of 3 were prepared in triplicate at 18 hpi and immobilized on RFP-TRAP beads (Chromotek, Rosemont, IL) according to manufacturer’s instructions. Washed beads were boiled in 2X SDS-PAGE loading buffer to elute the bound proteins, immunoprecipitated proteins were electrophoresed 5 mm into a regular 10% SDS-PAGE gel, stained with SimplyBlue SafeStain (Invitrogen, Carlsbad, CA) according to manufacturer’s instructions, and then 5 mm X 5 mm slices from the gel were excised. Orbitrap/iontrap tandem mass spectrometry on proteins eluted from gel slices and digested with trypsin was performed by the Southern Alberta Mass Spectrometry Facility at the University of Calgary.

### Mass spectrometry data analysis

Tandem mass spectra data were searched against a human protein database containing WT HSV proteins using Mascot version 2.7.0 (Matrix Science, London, UK), assuming trypsin as the digestion enzyme. Mascot was searched with a fragment ion mass tolerance of 0.020 Da and a parent ion tolerance of 10.0 PPM. Scaffold version 5.3.3 (Proteome Software Inc., Portland OR) was used to validate peptide and protein identifications. Peptide identifications were accepted if they could be established at greater than 95.0% probability by the Peptide Prophet algorithm [87] with Scaffold delta-mass correction. Protein identifications were accepted if they could be established at greater than 95.0% probability and contained at least one identified peptide. Protein probabilities were assigned by the Protein Prophet algorithm [88]. Proteins that contained similar peptides and could not be differentiated based on MS/MS analysis alone were grouped to satisfy the principles of parsimony.

Previously published common BioID contaminants [89] were removed from each dataset and proteins identified that met the following criteria were further analyzed: 1) a ≥ 95% protein and peptide threshold, 2) a minimum number of ≥ 3 identified unique peptides, with the exception of serine-threonine-protein phosphatase PP1-alpha and - gamma catalytic subunits that were retained due to a high degree of shared peptides 3) ≥ 5% normalized percent coverage. Total spectral counts were normalized to the average spectral counts of endogenously biotinylated mitochondrial proteins (mitochondrial average): acetyl-CoA carboxylase 1, pyruvate carboxylase, mitochondrial, methylcrotonoyl-CoA carboxylase subunit alpha, and propionyl-CoA carboxylase alpha chain. For both no-biotin and biotin-treated samples, normalization factors for each replicate were determined by dividing the highest mitochondrial average by each replicate’s mitochondrial average, and these factors were applied to all spectral counts within each replicate. Normalized spectral counts were determined by averaging values across three biological replicates, and no-biotin averages were subtracted from the plus-biotin averages to generate the final normalized spectral counts. The average of the percent coverage of three biological replicates of the no-biotin control samples was subtracted from the average of the percent coverage of the plus-biotin experimental samples to determine the normalized percent coverage.

Identified peptides and proteins associated with pUL21mCh were validated using Scaffold version 5.3.3. Common contaminants such as keratins were removed, and proteins identified that met the following criteria were further analyzed: 1) a ≥ 95% protein and peptide threshold, 2) a minimum number of ≥ 3 identified unique peptides, and 3) ≥ 5% averaged percent coverage. Average spectral counts and average percent coverage were calculated across three biological replicates.

### Purification of virions

Virions were purified as previously described [90]. Briefly, three confluent 150 mm dishes of HaCaT cells were infected at an MOI of 0.01. At 72 hpi, medium was collected and triple-cleared by centrifugation at 1,000 rpm for 10 minutes at 4°C. Clarified supernatants were layered onto a 30% sucrose (wt/vol, in PBS) cushion and centrifuged in a Beckman SW28 rotor at 23,000 rpm for three hours at 4°C. Pelleted virions were incubated in 140 µL BioID lysis buffer (50 mM Tris pH 7.5, 150 mM NaCl, 0.4% SDS, 1% NP-40, 1.5 mM MgCl_2_, 1 mM EGTA) containing protease inhibitors and 250 units of benzonase nuclease for 15 minutes at room temperature, resuspended in 70 µL 3X SDS-PAGE loading buffer, and analyzed by western blot.

### Analysis of pUL21mT biotinylation profile within purified 21-mT virions

Virions were purified as described above. Pelleted virions were treated with 0.01% saponin (Sigma) or 0.1% NP-40 (Igepal; Sigma) and 50 µM exogenous biotin in 140 µL of PBS for one hour and subsequently treated with 250 units of benzonase nuclease for 15 minutes at room temperature. Following treatment, virions were resuspended in 70 µL of 3X SDS-PAGE loading buffer and analyzed by western blot.

### Fluorescence recovery after photobleaching analysis

HeLa cells growing on 35 mm glass-bottom dishes (MatTek) were transfected using the X-tremeGene HP DNA transfection reagent (Roche) following the manufacturer’s protocol. At 24 hours post-transfection, cells were mounted onto an Olympus FV1000 laser scanning confocal microscope and maintained at 37°C in a humidified 5% CO_2_ environment. EGFP was excited using a 488 nm laser line set at 5% power. Images (512 by 512 pixels) were obtained using a 60X 1.42 NA oil immersion objective and a digital zoom factor of 6. Images were collected at a rate of 0.33 frames per second. Four frames were collected before the regions bounded by the red rectangles were photobleached by repeated scanning with a 405 nm laser set at 100% power for 300 ms. Before, during, and after photobleaching, fluorescence intensity in bleached and unbleached control regions was measured using FV10 ASW 4.01 software and the data were exported into Microsoft Excel for graphical presentation.

### Generation of Metascape GO enrichment network, STRING functional network map, and Venn diagrams

Functional enrichment analysis using the GO Molecular Functions, GO Biological Processes, and GO Cellular Components was conducted using Metascape (https://metascape.org) [58] with the following criteria: minimum overlap of 3, 0.01 p-value cutoff, and 1.5 minimum enrichment. Figures 4A, 8A, and 10A show simplified GO enrichment networks, displaying the top 10 clusters. Protein-protein interaction networks were generated using STRING version 12.0 (https://string-db.org) [59, 60] using the following criteria: physical network type, confidence network edge, experiments and databases active interaction sources, medium confidence (0.400) minimum required interaction score, and unlimited number of interactors shown (query proteins only). Edges connecting the nodes represent reported associations, with edge thickness corresponding to interaction confidence (thicker edges indicate higher confidence). Halos surrounding each node reflect normalized or averaged percent coverage, with darker halos indicating higher percent coverage. Only proteins with a normalized or averaged protein coverage of ≥ 5% were included in these analyses, and disconnected nodes were excluded. Venn diagrams were generated using Venny 2.1. (https://bioinfogp.cnb.csic.es/tools/venny/).

### Statistical analysis

Graphical illustrations and statistical analyses were performed using GraphPad Prism version 10.4.2.

## Supporting information

Supplementary Data 1 (BioID 18hpi)

Supplementary Data 2 (BioID 2hpi)

Supplementary Data 3 (UL21mCh pulldown)

## Acknowledgements

This work was supported by the Canadian Institutes of Health Research (CIHR) operating grant 162162, CIHR Project Grant 186187, Natural Sciences and Engineering Research Council of Canada Discovery Grants RGPIN/04249 and RGPIN/06733 and Canada Foundation for Innovation (CFI) award 16389 to BWB. SMH was supported in part by an Ontario Graduate Scholarship and Robert Sutherland and R. Samuel McLaughlin Fellowships from Queen’s University. The funders had no role in study design, data collection and interpretation, or the decision to submit the work for publication. We thank Dr. Laurent Brechenmacher from the Southern Alberta Mass Spectrometry Centre at the University of Calgary for expert assistance with mass spectrometry. We are grateful to all members of the Banfield laboratory for helpful comments on the manuscript.

## Supplementary Figure Legends

**Supplementary Figure 1.**
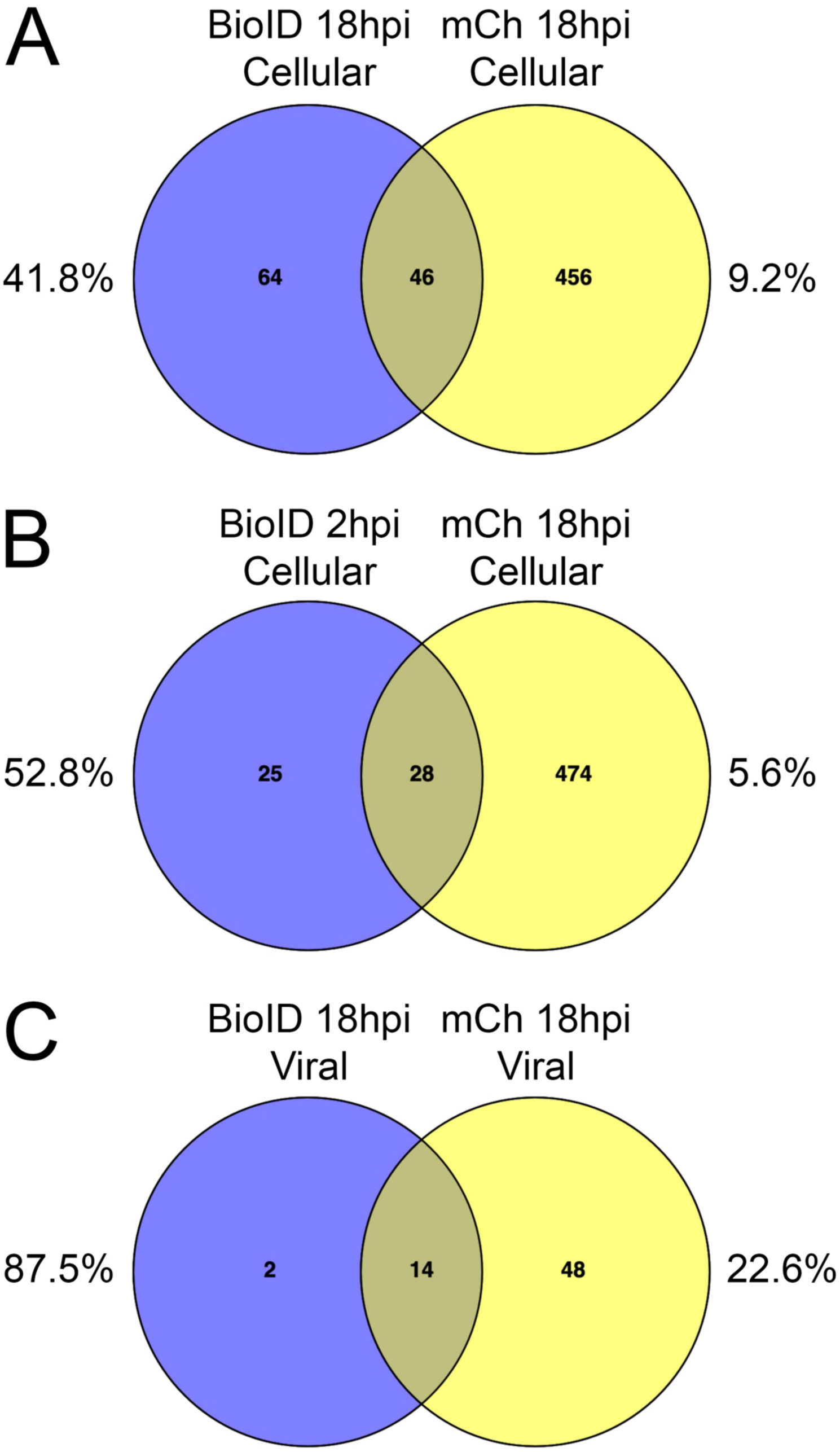
Comparisons of identified proteins in proximity to pUL21mT and interacting with pUL21mCh at 18 hours post-infection. **(A)** Venn diagram of the number of unique and common cellular proteins in proximity to pUL21mT and interacting with pUL21mCh at 18 hours post-infection (hpi). **(B)** Venn diagram of the number of unique and common cellular proteins in proximity to tegument-delivered pUL21mT at 2 hpi and interacting with pUL21mCh at 18 hpi. **(C)** Venn diagram of the number of unique and common viral proteins in proximity to pUL21mT and interacting with pUL21mCh at 18 hpi. Percentages in each panel reflect the proportion of proteins in each dataset that fall within the intersection.

**Supplementary Figure 2.**
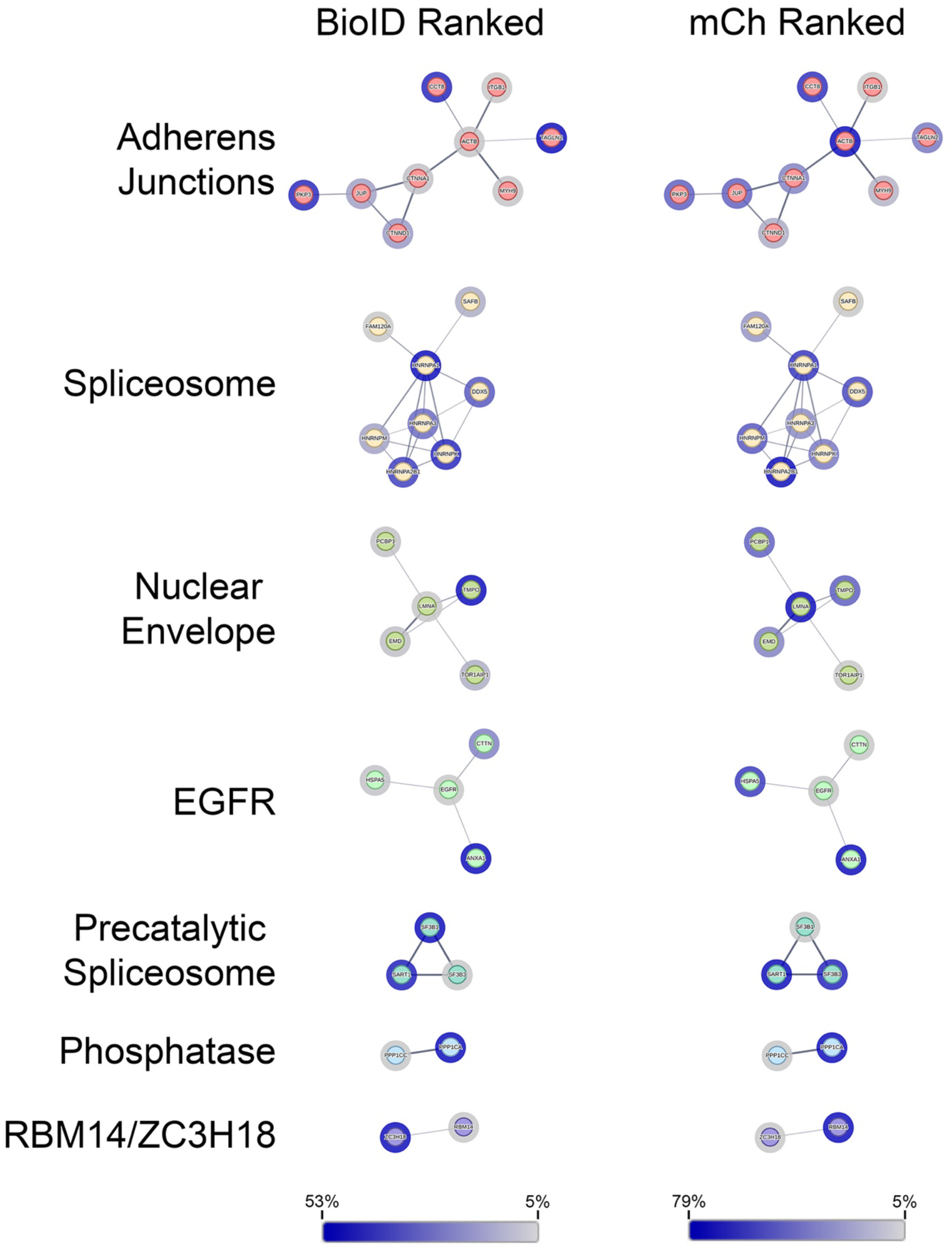
STRING analysis of commonly identified cellular proteins in proximity to pUL21mT and interacting with pUL21mCh at 18 hours post-infection. Identified cellular proteins were analyzed, clustered, and colour-coded using the STRING database to visualize known protein-protein interactions. The protein-protein interaction network of each individual cluster is shown. EGFR (epidermal growth factor receptor). Edges connecting the nodes represent reported associations, with edge thickness corresponding to interaction confidence (thicker edges indicate higher confidence). Halos surrounding each node reflect percent coverage, with darker halos indicating higher percent coverage. Only proteins with a normalized or averaged protein coverage of ≥ 5% were included in this analysis, and disconnected nodes are not shown.

## Supplementary Tables

**Supplementary Table 1.**
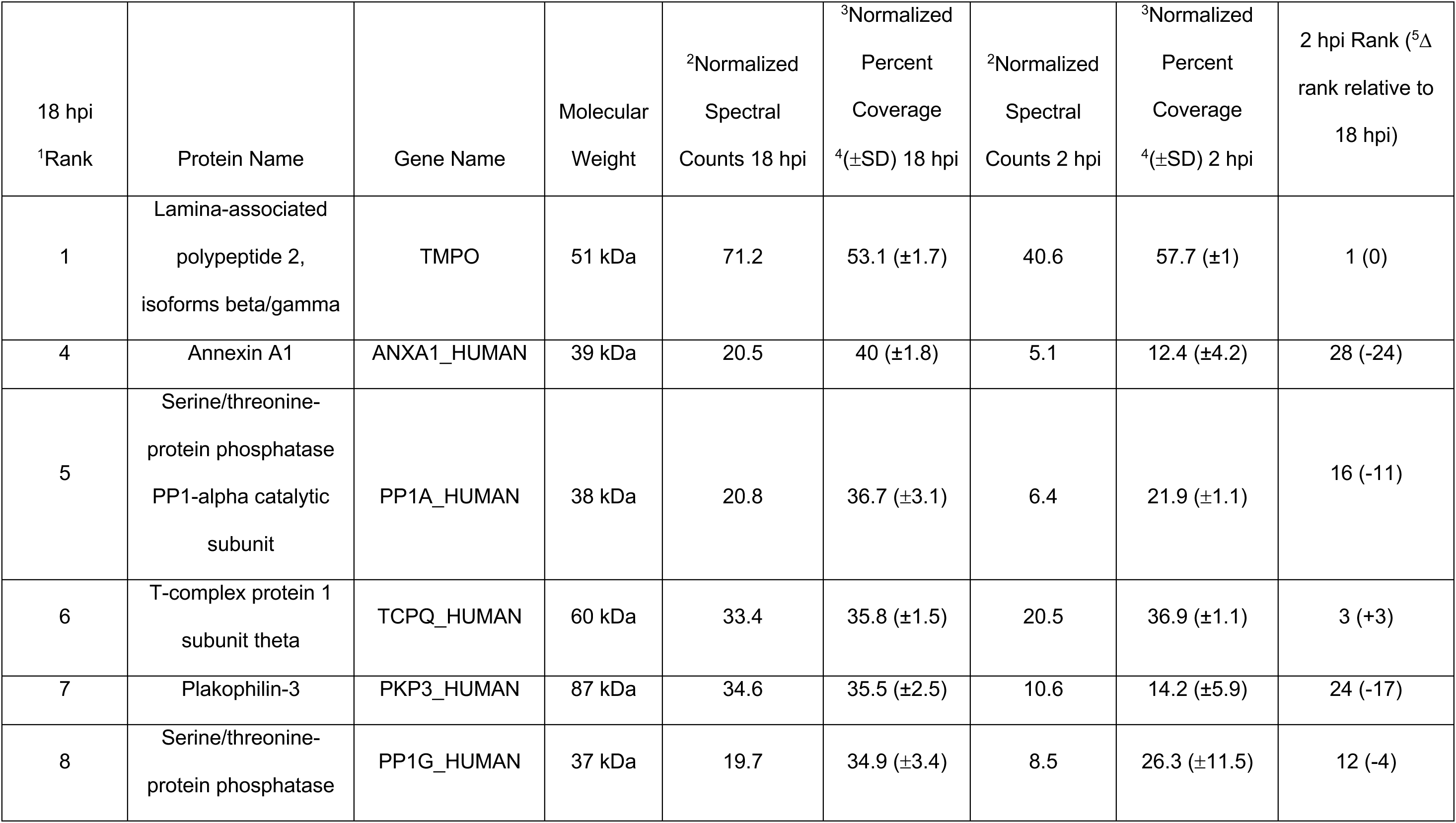
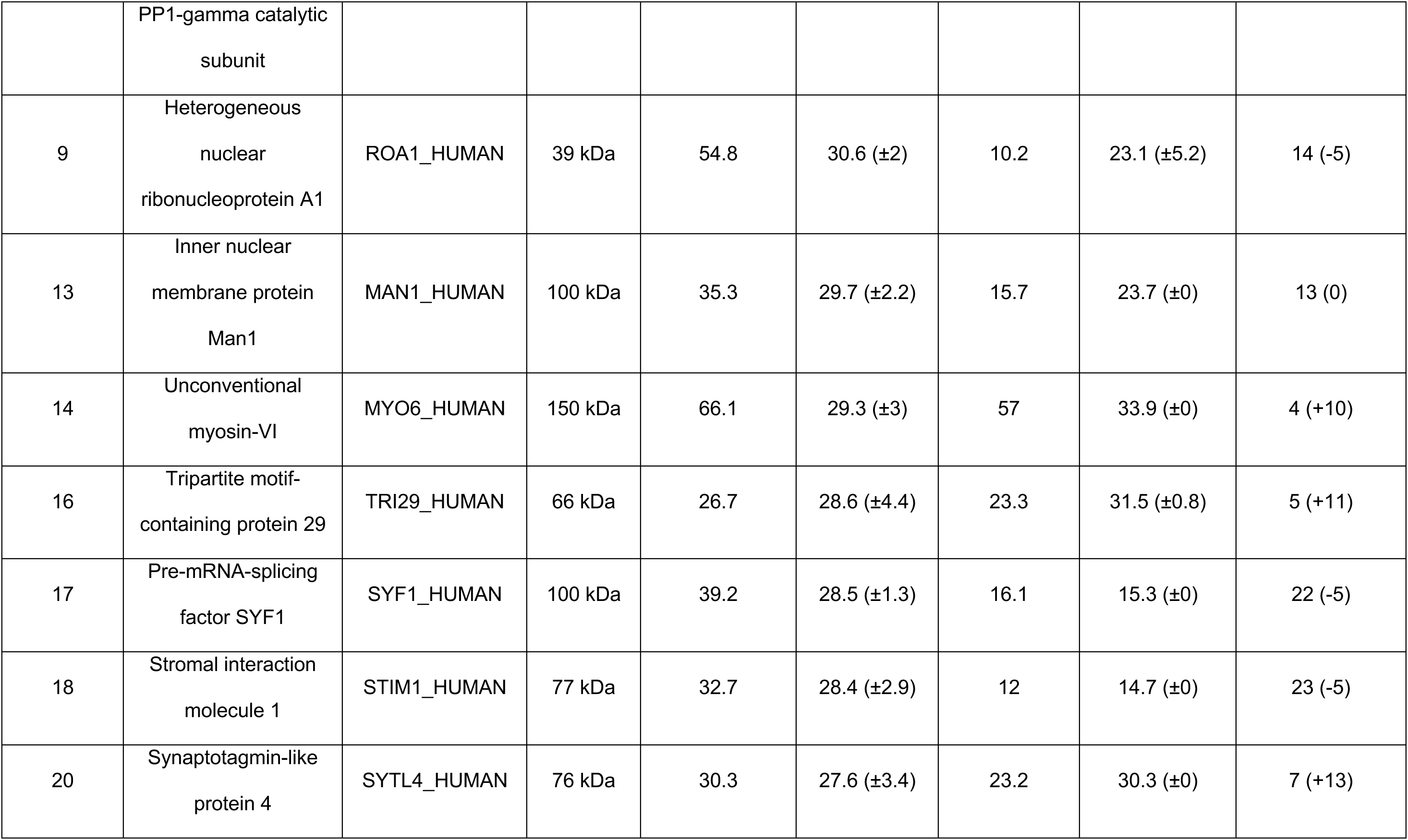
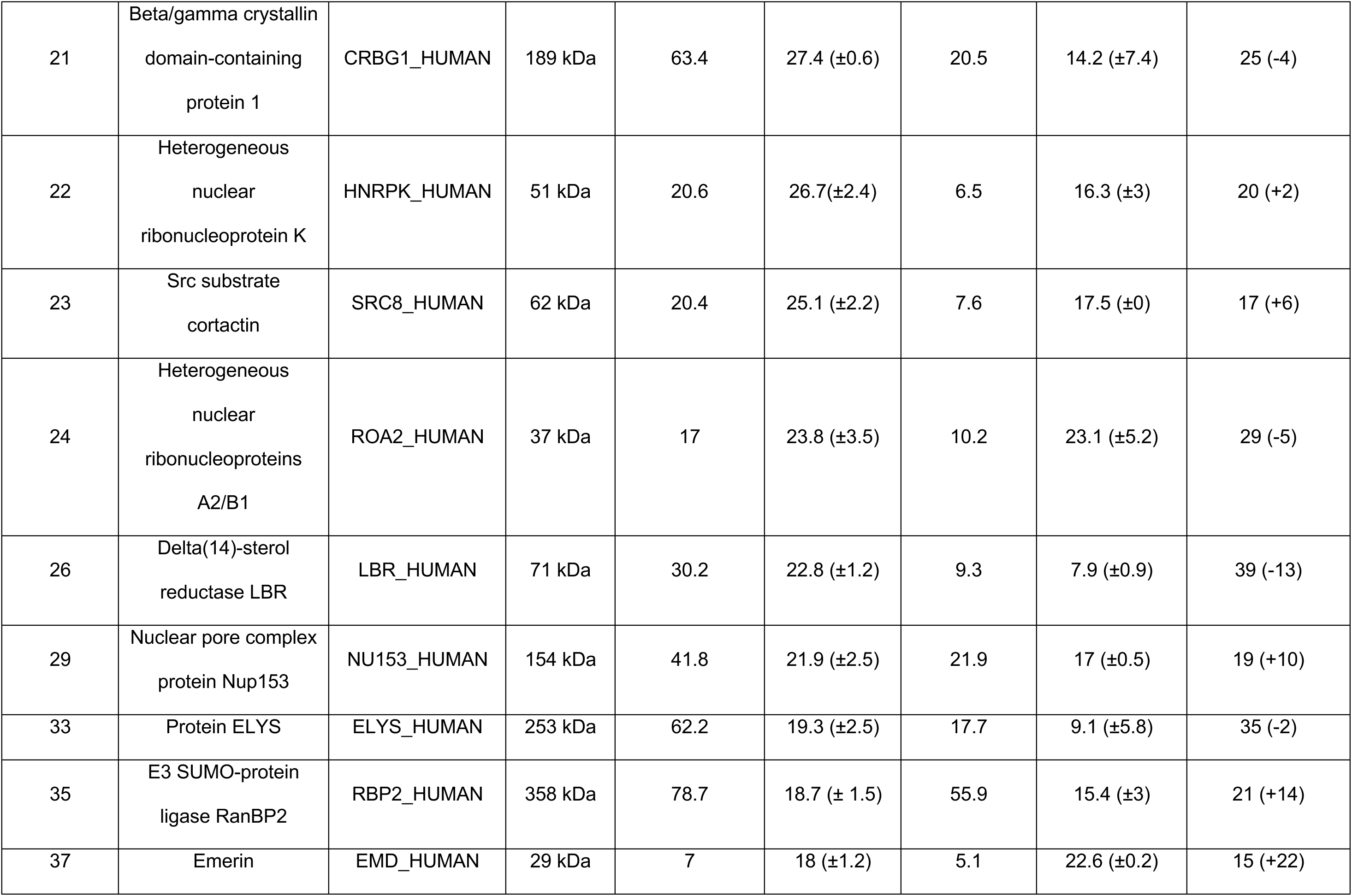
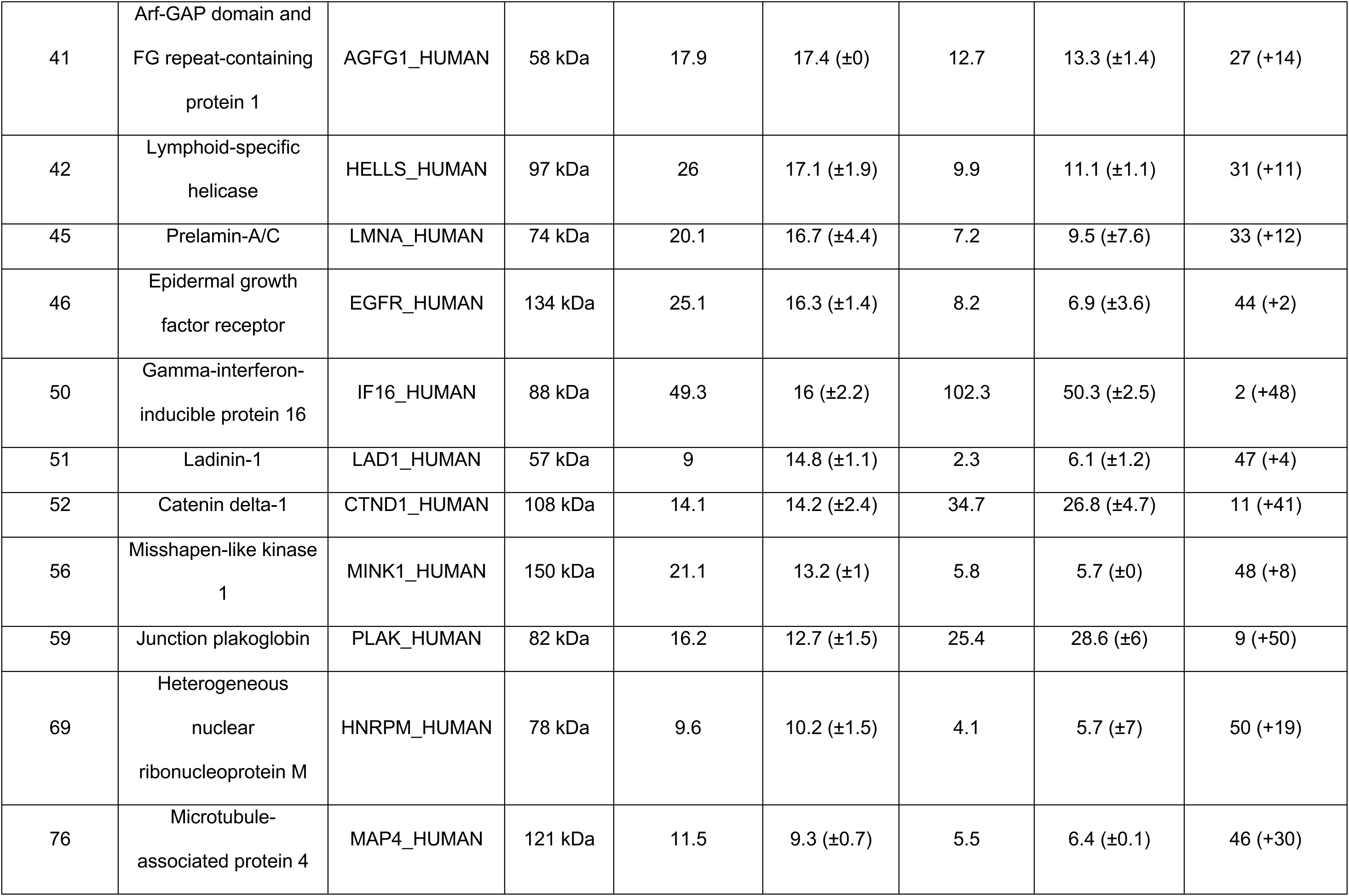
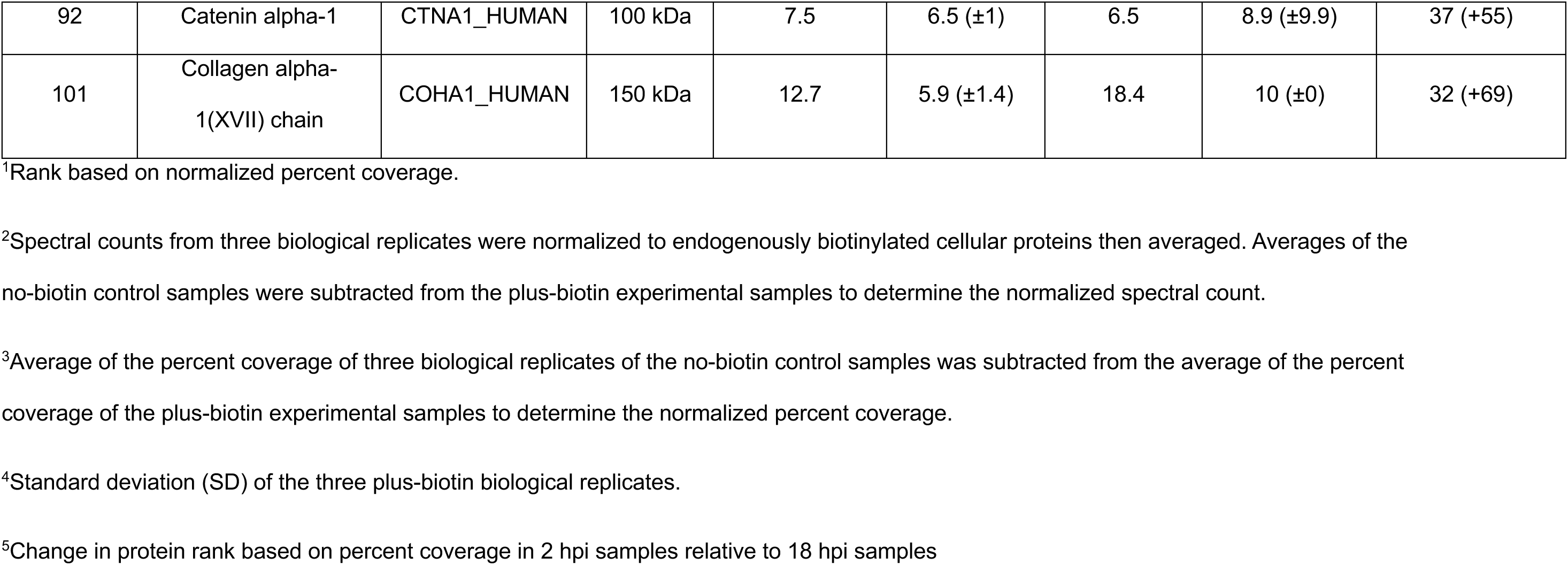
Common Proteins in Proximity to pUL21mT at 18 hpi and 2 hpi.

**Supplementary Table 2.**
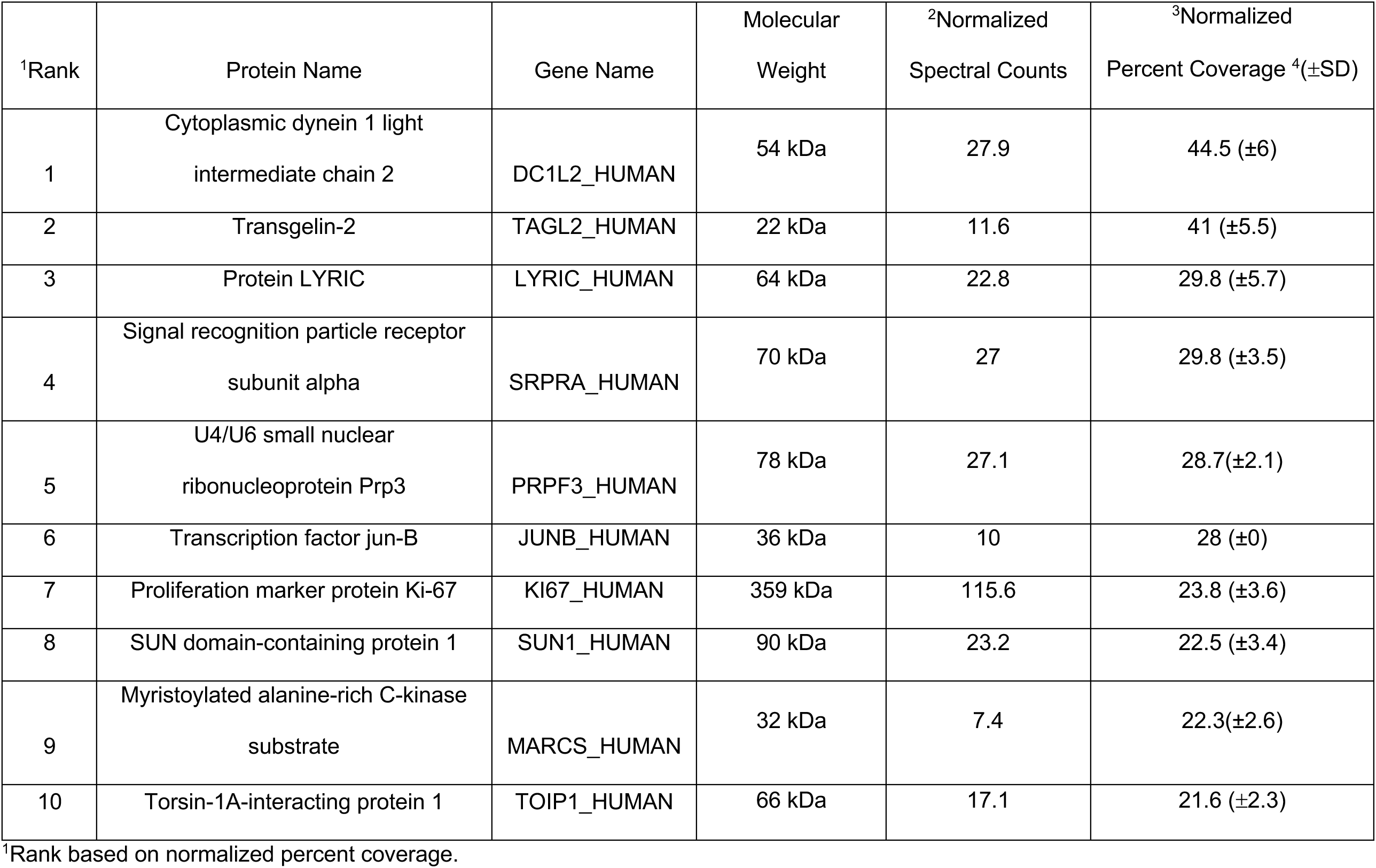
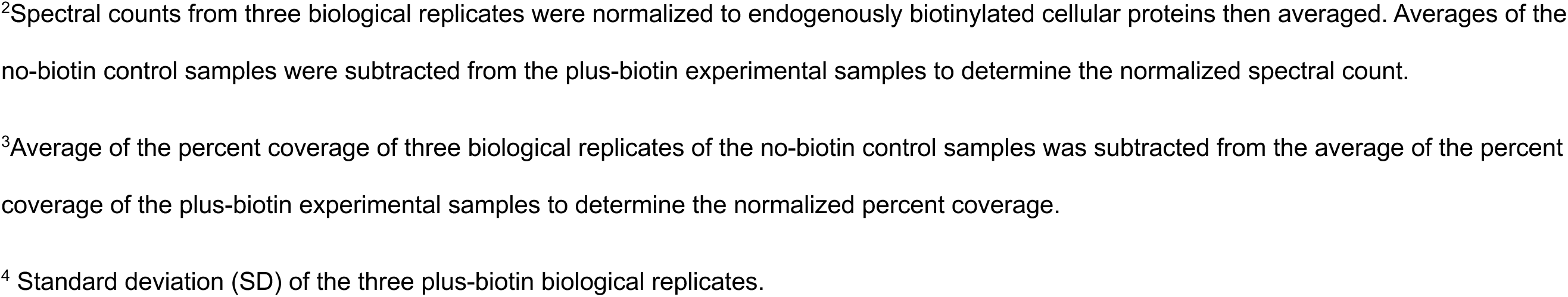
Top 10 Unique Proteins in Proximity to pUL21mT at 18 hpi.

| <sup>1</sup>Rank | Protein Name | Gene Name | Molecular Weight | <sup>2</sup>Normalized Spectral Counts | <sup>3</sup>Normalized Percent Coverage <sup>4</sup>(±SD) |
| --- | --- | --- | --- | --- | --- |
| 1 | Cytoplasmic dynein 1 light intermediate chain 2 | DC1L2_HUMAN | 54 kDa | 27.9 | 44.5 (±6) |
| 2 | Transgelin-2 | TAGL2_HUMAN | 22 kDa | 11.6 | 41 (±5.5) |
| 3 | Protein LYRIC | LYRIC_HUMAN | 64 kDa | 22.8 | 29.8 (±5.7) |
| 4 | Signal recognition particle receptor subunit alpha | SRPRA_HUMAN | 70 kDa | 27 | 29.8 (±3.5) |
| 5 | U4/U6 small nuclear ribonucleoprotein Prp3 | PRPF3_HUMAN | 78 kDa | 27.1 | 28.7(±2.1) |
| 6 | Transcription factor jun-B | JUNB_HUMAN | 36 kDa | 10 | 28 (±0) |
| 7 | Proliferation marker protein Ki-67 | KI67_HUMAN | 359 kDa | 115.6 | 23.8 (±3.6) |
| 8 | SUN domain-containing protein 1 | SUN1_HUMAN | 90 kDa | 23.2 | 22.5 (±3.4) |
| 9 | Myristoylated alanine-rich C-kinase substrate | MARCS_HUMAN | 32 kDa | 7.4 | 22.3(±2.6) |
| 10 | Torsin-1A-interacting protein 1 | TOIP1_HUMAN | 66 kDa | 17.1 | 21.6 (±2.3) |

<sup>1</sup>Rank based on normalized percent coverage.
<sup>2</sup>Spectral counts from three biological replicates were normalized to endogenously biotinylated cellular proteins then averaged. Averages of the no-biotin control samples were subtracted from the plus-biotin experimental samples to determine the normalized spectral count.
<sup>3</sup>Average of the percent coverage of three biological replicates of the no-biotin control samples was subtracted from the average of the percent coverage of the plus-biotin experimental samples to determine the normalized percent coverage.
<sup>4</sup> Standard deviation (SD) of the three plus-biotin biological replicates.

**Supplementary Table 3.** Top 10 Unique Proteins in Proximity to pUL21mT at 2.

| <sup>1</sup>Rank | Protein Name | Gene Name | Molecular Weight | <sup>2</sup>Normalized Spectral Counts | <sup>3</sup>Normalized Percent Coverage <sup>4</sup>(±SD) |
| --- | --- | --- | --- | --- | --- |
| 1 | Thioredoxin | THIO_HUMAN | 12 kDa | 7.2 | 31.4 (±0.5) |
| 2 | Erbin | ERBIN_HUMAN | 158 kDa | 37.8 | 29.3 (±7.8) |
| 3 | Pleckstrin homology domain-containing family A member 5 | PKHA5_HUMAN | 127 kDa | 33.8 | 28.1 (±0.7) |
| 4 | Myosin light polypeptide 6 | MYL6_HUMAN | 17 kDa | 2.4 | 17.2 (±0.3) |
| 5 | Sickle tail protein homolog | SKT_HUMAN | 214 kDa | 26.9 | 14 (±0) |
| 6 | Transmembrane protein 263 | TM263_HUMAN | 12 kDa | 2.4 | 11.8 (±1.1) |
| 7 | Plakophilin-1 | PKP1_HUMAN | 83 kDa | 5.5 | 9.5 (±0) |
| 8 | Rho GTPase-activating protein 21 | RHG21_HUMAN | 217 kDa | 15.1 | 8.9 (±1.9) |
| 9 | Protein FAM83H | FA83H_HUMAN | 127 kDa | 7.9 | 8.6 (±0) |
| 10 | Tubulin beta-4B chain | TBB4B_HUMAN | 50 kDa | 2.8 | 7.5 (±0) |

<sup>1</sup>Rank based on normalized percent coverage.
<sup>2</sup>Spectral counts from three biological replicates were normalized to endogenously biotinylated cellular proteins then averaged. Averages of the no-biotin control samples were subtracted from the plus-biotin experimental samples to determine the normalized spectral count.
<sup>3</sup>Average of the percent coverage of three biological replicates of the no-biotin control samples was subtracted from the average of the percent coverage of the plus-biotin experimental samples to determine the normalized percent coverage.
<sup>4</sup> Standard deviation (SD) of the three plus-biotin biological replicates.

**Supplementary Table 4.** Common Cellular Proteins Identified by BioID at 18 hpi and Affinity-Purified with pUL21mCh at 18 hpi.

| <sup>1</sup>Rank | Gene Name |
| --- | --- |
| 1 | TMPO |
| 2 | TAGL2_HUMAN |
| 3 | ANXA1_HUMAN |
| 4 | PP1A_HUMAN |
| 5 | TCPQ_HUMAN |
| 6 | PKP3_HUMAN |
| 7 | PP1G_HUMAN |
| 8 | ROA1_HUMAN |
| 9 | LYRIC_HUMAN |
| 10 | SRPRA_HUMAN |
| 11 | TRI29_HUMAN |
| 12 | HNRPK_HUMAN |
| 13 | SRC8_HUMAN |
| 14 | ROA2_HUMAN |
| 15 | MARCS_HUMAN |
| 16 | TOIP1_HUMAN |
| 17 | EMD_HUMAN |
| 18 | ROA3_HUMAN |
| 19 | PCBP1_HUMAN |
| 20 | SF3B1_HUMAN |
| 21 | BIP_HUMAN |
| 22 | DDX3X_HUMAN |
| 23 | LMNA_HUMAN |
| 24 | EGFR_HUMAN |
| 25 | SNUT1_HUMAN |
| 26 | LAD1_HUMAN |
| 27 | CTND1_HUMAN |
| 28 | DDX18_HUMAN |
| 29 | PLAK_HUMAN |
| 30 | ZCH18_HUMAN |
| 31 | PAI2_HUMAN |
| 32 | TR150_HUMAN |
| 33 | HNRPM_HUMAN |
| 34 | RBM14_HUMAN |
| 35 | SAFB1_HUMAN |
| 36 | NAT10_HUMAN |
| 37 | FAS_HUMAN |
| 38 | PEPL_HUMAN |
| 39 | CTNA1_HUMAN |
| 40 | NUCL_HUMAN |
| 41 | SF3B3_HUMAN |
| 42 | COHA1_HUMAN |
| 43 | MYH9_HUMAN |
| 44 | ADNP_HUMAN |
| 45 | ITB1_HUMAN |
| 46 | F120A_HUMAN |

<sup>1</sup>Proteins ranked in order of normalized percent coverage obtained in 18 hpi BioID experiment.

**Supplementary Table 5.**
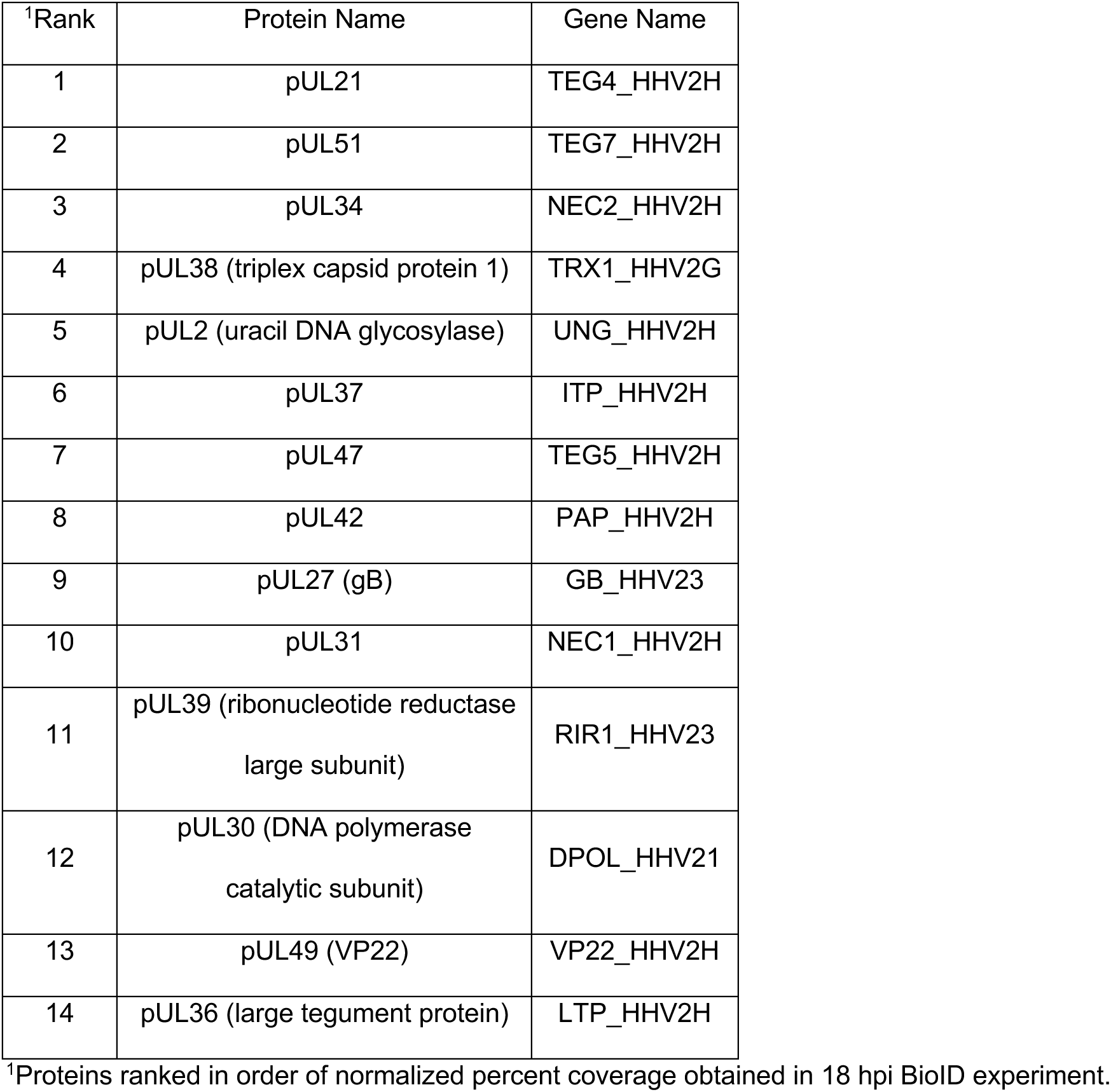
Common Viral Proteins Identified by BioID at 18 hpi and Affinity-Purified with pUL21mCh at 18 hpi.

**Supplementary Table 6.** Common Cellular Proteins Identified by BioID at 2 hpi and Affinity-Purified with pUL21mCh.

| <sup>1</sup>Rank | Gene Name |
| --- | --- |
| 1 | TMPO |
| 2 | TCPQ_HUMAN |
| 3 | TRI29_HUMAN |
| 4 | PLAK_HUMAN |
| 5 | CTND1_HUMAN |
| 6 | PP1G_HUMAN |
| 7 | ROA1_HUMAN |
| 8 | EMD_HUMAN |
| 9 | PP1A_HUMAN |
| 10 | SRC8_HUMAN |
| 11 | MYL6_HUMAN |
| 12 | HNRPK_HUMAN |
| 13 | PKP3_HUMAN |
| 14 | ANXA1_HUMAN |
| 15 | ROA2_HUMAN |
| 16 | COHA1_HUMAN |
| 17 | LMNA_HUMAN |
| 18 | CTNA1_HUMAN |
| 19 | FA83H_HUMAN |
| 20 | TBB4B_HUMAN |
| 21 | CAV1_HUMAN |
| 22 | EGFR_HUMAN |
| 23 | ANXA2_HUMAN |
| 24 | LAD1_HUMAN |
| 25 | EF1A1_HUMAN |
| 26 | HNRPM_HUMAN |
| 27 | SPB5_HUMAN |
| 28 | CAZA1_HUMAN |

<sup>1</sup>Proteins ranked in order of normalized percent coverage obtained in 2 hpi BioID experiment.

